# Human induced pluripotent stem cell-derived vessels as dynamic atherosclerosis model on a chip

**DOI:** 10.1101/2020.11.27.401034

**Authors:** A. Mallone, C. Gericke, V. Hosseini, K. Chahbi, W. Haenseler, M. Y. Emmert, A. von Eckardstein, J. H. Walther, V. Vogel, B Weber, S. P. Hoerstrup

## Abstract

Atherosclerosis is an arterial disease characterized by intravascular plaques. Disease hallmarks are vessel stenosis and hyperplasia, eventually escalating into plaque rupture and acute clinical presentations. Innate immune cells and local variations in hemodynamics are core players in the pathology, but their mutual relationship has never been investigated before due to the lack of modeling systems with adequate degree of complexity. Here, we combined computational fluid dynamics and tissue-engineering to achieve, for the first time *in vitro*, full atherosclerotic plaque development. Our model incorporates induced pluripotent stem cell-derived populations into small-caliber arteries that are cultured in atheroprone conditions. Using machine-learning-aided immunophenotyping, molecular and nanoprobe-based tensile analyses, we found that immune cells, extracellular matrix components and tensional state were comparable between *in vitro* and *ex vivo* human lesions. Our results provide further insights into the relation between hemodynamics and inflammation, introducing a versatile, scalable modeling tool to study atherosclerosis onset and progression.

## Introduction

Atherosclerosis is an inflammatory disease of the arterial walls associated with life-threatening events such as myocardial infarction or stroke^1,2^. Disease hallmarks are fibrotic and fatty intra-vessel plaques, which cause abnormal vascular narrowing^3^. Plaque erosion followed by thrombus formation can lead to diverse outcomes depending on the downstream affected organ or vasculature^4^. Besides cases of ascertained genetic predisposition (i.e., familial hypercholesterolemia), multiple other factors can contribute to sporadic atherosclerosis. Among those, dyslipidemia with high blood levels of low-density lipoproteins (LDL) is the main risk factor, followed by chronic inflammation as well as age- and lifestyle-related comorbidities such as diabetes and hypertension^5–8^. Some of these conditions have been associated with systemic changes in blood hemodynamics, local alterations of the flow pattern (e.g., vortexes and turbulences), and mechanical stress on the vessel walls^9–11^. As a result, flow variations promote local sub-luminal accumulation of LDL, providing the initial trigger for an inflammatory response^12–16^.

*In vitro* models based on 2D-adherent cell cultures allowed a partial understanding of the activation mechanisms of endothelial, smooth muscle, and immune cells upon changes in flow dynamics and lipoproteins concentration^8,17,18^. Some 3D tissue-engineered tools (i.e., microtissues and organoids) attempted to mimic early and late stages of the disease^19–21^ while animal models provided insights on plaque extracellular matrix (ECM) remodeling by inflammatory cells^22–25^. However, to the best of our knowledge, none of the above has been able to recapitulate the complex process of plaque build-up and ECM remodeling in a tissue-engineered replica of a human artery. Furthermore, it remains largely unknown how immune cells behave in regions of hemodynamic perturbation within a human, native-like vascular environment. Here, we hypothesized that it is possible to predict the location of regions of LDL accumulation regions in a vessel using *in silico* modeling, thus allowing for accurate prediction of potential plaque deposition areas. We further hypothesized that it is possible to accelerate plaque development *in vitro*, a long-lasting process in humans^1^, by introducing pathophysiological levels of circulating inflammatory cells, LDL, and altered flow patterns. In this work, we developed computational fluid dynamic (CFD) simulation of atherosclerotic plaque seeding in tissue-engineered vessels. This model allowed us to predict preferential intravascular hot spots of LDL accumulation and to measure the sub-endothelial, potentially LDL-caused hindrance. To validate our predictions, we replicated such an environment *in vitro* by setting up a customized fluidic device hosting multiple 3D tissue-engineered small-caliber vascular grafts. The vessels were generated from human induced pluripotent stem cell (iPSC) – derived cells and were used to model atherosclerosis through daily exposure to atheroprone stimuli. Our work pioneers multiplexing approaches in vascular tissue-engineering and contributes to the optimization efforts in atherosclerosis modeling, serving as a starting point for other in-depth analyses focused on more specific stages of disease progression.

## Results

### 1. iPSC-derived 3D human vessels display tissue architecture comparable to human native vasculature

To develop small-caliber arterial vessels, we first differentiated iPSCs into arterial endothelial cells (iECs) (Figure 1a-d, Supplementary Fig.1). We characterized the iECs using flow-cytometry and analyzed the flow data by unbiased machine-learning-guided approaches (here: t-distributed stochastic neighbor embedding, tSNE) and data-driven clustering algorithms (Figure 1b). We observed that iECs mainly consisted of CD31+/CD144+/CD68+^high^ cells (cluster 1, 92.45 %, Figure 1b, Supplementary Fig.1). The iECs phenotype was further confirmed via immunofluorescence analysis (IF) and by investigating their gene expression profile (Figure 1c, Supplementary Fig. 1). iECs had no significant differences in expression levels of the key endothelial genes *VE-CADH* and *CD31* when compared to control endothelial cell lines such as human umbilical vein endothelial cells (HUVECs; *VE-CADH p* = 0.45, *CD31 p* = 0.09) and human brain microvascular endothelial cells (HBMECs; *VE-CADH p* = 0.39, *CD31 p* = 0.07). To test iECs functionality, we performed a tube formation assay, showing that iECs could sprout and generate new micro-vascular networks (Figure 1d). Next, we differentiated contractile smooth muscle cells (iSMCs) from syngeneic iPSCs (Figure 1e-k, Supplementary Fig. 2). In IF experiments (Figure 1f-g, Supplementary Fig. 2), iSMCs displayed typical smooth muscle markers such as smooth muscle myosin heavy chain (SMMHC), transgelin (SM22α) and alpha actin (αSMA). We further confirmed the SMCs phenotype by looking at the gene expression profile (Figure 1d) where we observed the upregulation of the signature smooth muscle genes *aSMA* (*p* = 0.03) and *SMTN* (*p* = 0.03). We investigated iSMCs functionality using a hydrogel contraction and a wound healing assay (Figure 1h-j). We confirmed the iSMCs contractile phenotype showing a behavior of iSMCs comparable to human umbilical vein myofibroblasts (HUVMs vs iSMCs ns: *p* > 0.05). With the wound healing assay we further assessed the ability of iSMCs to convert to a disease-associated synthetic state and acquire migration skills, as described in literature^17^.

**Figure 1.**
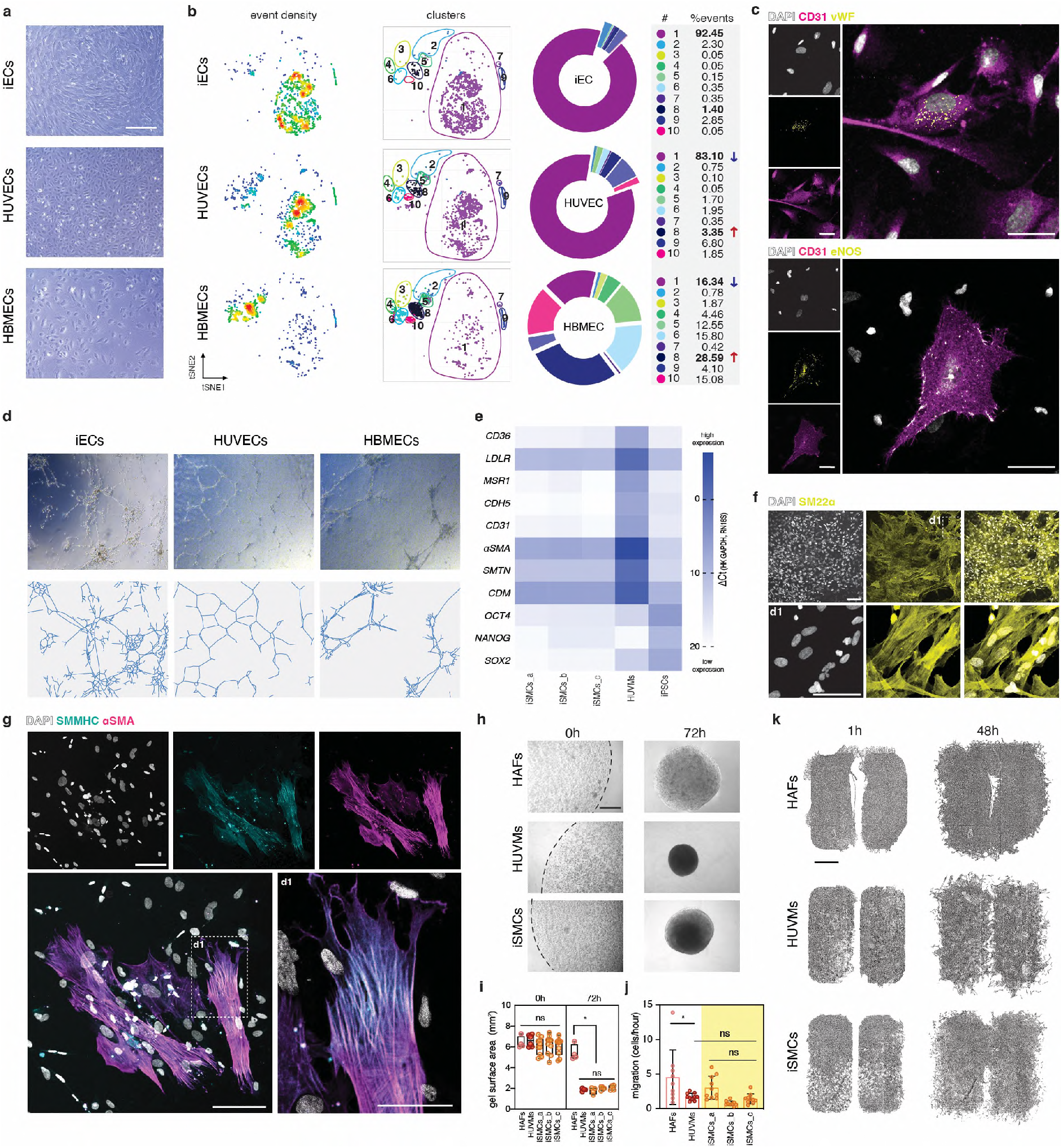
Generation and characterization of syngeneic endothelial and smooth muscle cells from iPSCs for vascular tissue engineering. (a) Bright field images of iECs, HUVECs and HBMECs. Scale bar 250 μm. (b) Flow-cytometry characterization of different endothelial cells (iECs, HUVECs, and HBMECs) with a multicolor and poli-marker panel. Multidimensional data are showed in 2D tSNE plots. Event density, FlowSOM clusters and cluster abundances (% of events) are shown. Red arrows indicate cell clusters with increased abundance in HUVECs and HBMECs (vs iECs), and blue arrows indicate cell clusters with decreased abundance in HUVECs and HBMECs (vs iECs). (c) IF characterization of iECs. iECs express platelet endothelial cell adhesion molecule precursor (CD31), von Willebrand factor (vWF) and endothelial nitric oxide synthase (eNOS). Scale bars 50μm. (d) Characterization of iECs functionality via tube formation assay. (e) Characterization of iSMC differentiation through RT-qPCR. HUVMs and iPSCs are used as positive and negative controls, respectively. (f) iSMCs expression of the smooth muscle protein transgelin (SM22α). d1= detail. Scale bars 100μm. (g) iSMCs expression of smooth muscle myosin heavy chain (SMMHC) and smooth muscle alpha actin (αSMA). d1= detail. Scale bars 100μm. (h-i) Characterization of iSMCs functionality using a fibrin gel contraction assay. Assay duration = 72h. Scale bar 250μm. (j-k) Characterization of iSMCs plasticity with wound healing assay. Assay duration = 48h, Scale bar 2mm. *p* values are reported in the text.

To generate native-like human vessels, we developed a versatile tissue-engineering approach based on a custom-made fluidic device. The device was connected to an automated, pulsatile flow system creating a dynamic and controlled culturing environment (Figure 2a-e, Supplementary Fig. 3, Supplementary Video1-2, Supplementary File1-5). To understand the inner flow distribution, we developed a CFD model of the fluid-domain considering perfusion through the inlet pipe at a constant flow rate, fluid density, and dynamic viscosity (Supplementary Fig. 4-5-6). We obtained a stable steady-state solution with low Reynolds numbers, suggesting the presence of a laminar flow. The model predicted arterial-like flow velocity within the vessels, ranging between 0.03 m/s at the extremities and 0.15 m/s in the lumen, and a total pressure ranging from 127 Pa at the inlets to about 20 Pa at the outlets (Figure 2c-d, Supplementary Fig. 4-5). Importantly, the results were comparable at each position for all the vessels, thus suggesting the suitability of the device for multiplexing and high-reproducibility approaches (Figure 2c-d, Supplementary Fig. 3, Supplementary File6-7). We then combined iECs and iSMCs to assemble tissue-engineered vessels from human induced pluripotent stem cells (hiTEVs), cultured them within the fluidic device, and compared them to human explants. hiTEVs showed lumen endothelialization concomitant with collagen accumulation throughout the para-luminal area, and the formation of confined, multicellular media layers perpendicularly aligned to the flow direction (Figure 2f-h). Taken together, these observations suggest that our culturing setup and tissue-engineering strategy promote the assembly of vessels with a stratified, native-like microanatomy.

**Figure 2.**
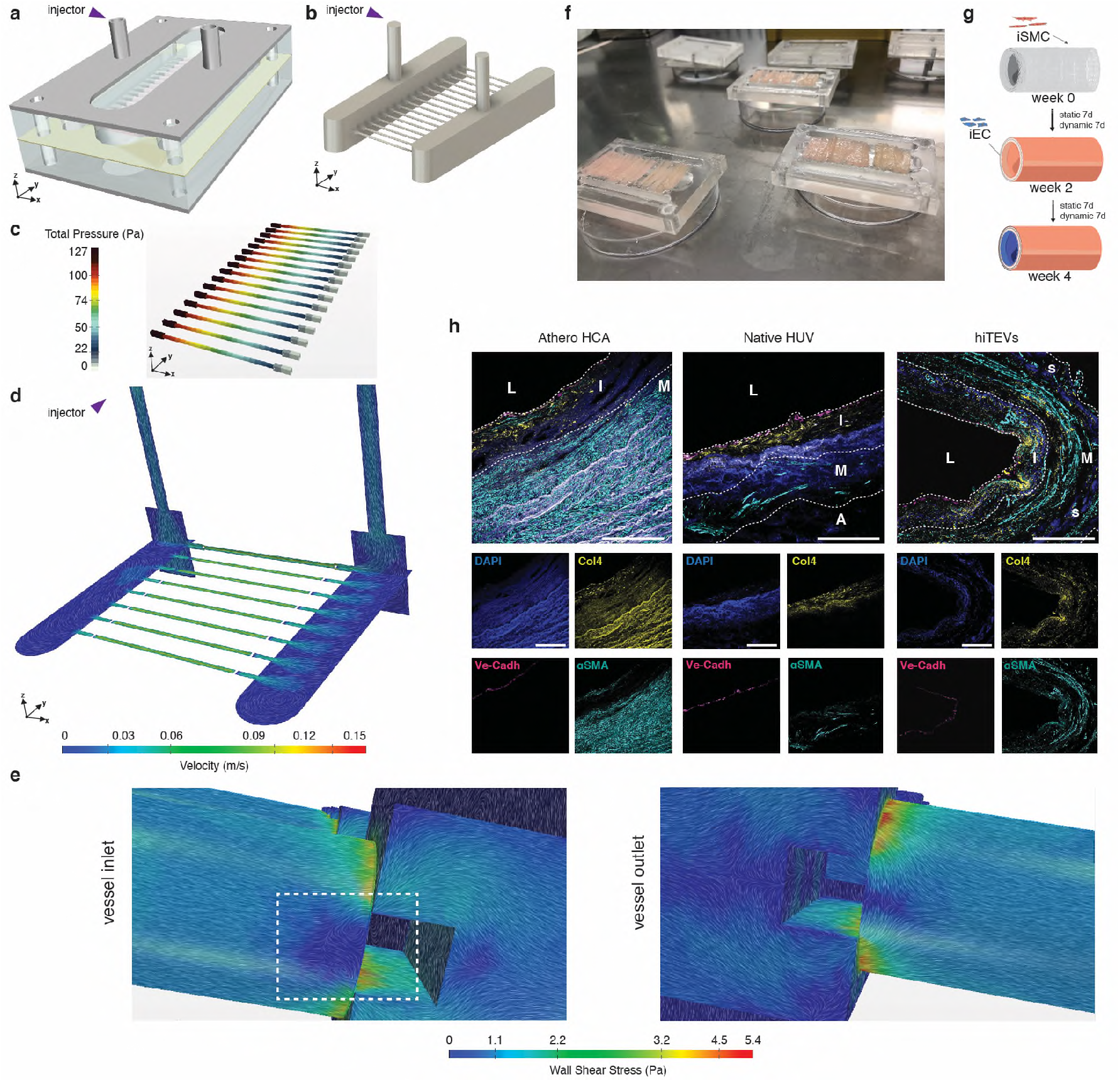
Small-caliber human iPSC-derived tissue engineered vessels. (a) CAD model of the assembled fluidic chamber. (b) Flow domain. Inner volume of the chamber occupied by the culturing medium. (c) Depiction of the pressure gradient within the tissue engineered vessels (hiTEVs). (d) Cross-sectional and longitudinal view of the flow velocity in the chamber and in the hiTEVs. (e) Wall shear stress at the hiTEV-inlets and hiTEV-outlets. The velocity field is superimposed to the flow domain and depicted using line-integral convolution. Whirlpools are highlighted in the rectangle. (f) Multiple, sterile fluidic chambers loaded with hiTEVs after intraluminal seeding of iECs. (g) Schematics of the tissue-engineering strategy adopted in the study. (h) IF comparison of carotid arteries from patients’ samples (Athero HCA), Native human umbilical vessels (Native HUV), and tissue-engineered arteries from iPSCs (hiTEVs). L = lumen, I = intima layer, M = media layer, S = PGA/P4HB scaffold remains. Scale bars 100μm.

### 2. Computational fluid dynamic analysis predicts atheroprone, hyperplasic, LDL-laden inflammatory niches in tissue-engineered small-caliber vessels cultured *in vitro*

To develop an *in silico* model of atherosclerosis we adapted our CFD simulation by introducing LDL particles at the inlet boundary as a concertation field (passive scalar). We modeled advection, drag, lift forces and random motion of LDL particles diffusing in the fluid domain (Figure 2e, Figure b-g). We elaborated a dynamic surface growth prediction where we extrapolated the putative degree of LDL deposition in the intima (growth rate) (Figure 3c-d). Importantly, we identified possible athero-susceptible regions at each vessel inlet. The flow conditions in such regions were absent elsewhere and were characterized by low wall shear stress (WSS) ranging from 0.2 to 0.4 Pa, by the presence of whirlpools and steady vortices, and by a predilection for impacting and accumulating LDL (Figure 3d-f). To validate our *in silico* data, we developed an *in vitro* disease model using hiTEVs (Figure 3k). We regularly perfused hiTEVs with LDL at hyperlipidemic concentrations and with inflammatory cells. Inflammatory cells consisted of macrophage precursors (iMPs) originated from iPSCs (Figure 3j, Supplementary Fig. 7). The iMPs upregulated monocyte-macrophage lineage-specific genes when compared to undifferentiated hiPSCs (CD36 *p* = 0.03, MSR1 *p* = 0.03), similarly to THP-1 monocytes/macrophages (Supplementary Fig. 7). A flow-cytometry characterization revealed that iMPs comprise three sub-populations: pro-inflammatory monocytes/macrophages expressing CD14^high^/CD16^high^/CD11b^high^ (Mo_pro-inf_1, 14.2% ± 6.8, and Mo_pro-inf_2, 42.2% ± 9.1), and macrophages being CD14^low^/CD16^high^/CD11b^high^ (Mf 1, 40.8% ± 12.6) (Supplementary Fig. 7). Upon differentiation, iMP-derived macrophages adhered to the culture flask, developed a large CD11b+ cell body, and could uptake LDL in a dose-dependent manner (Figure 3h-j, Supplementary Video3). At the end of four weeks of modeling time, we observed the formation of localized bulges at the hiTEV-inlets, similar to atherosclerotic plaques (Figure 3l). The hiTEV-outlets were not affected by tissue hyperplasia and were hereafter chosen as internal negative controls of plaque deposition. We then investigated whether such plaque-like structures included CD11b+ macrophages differentiating from the perfused iMPs. We found that iMPs in the pulsatile stream adhered to the vessel lumen, and migrated into the intima layer, gathering in CD11b+ intra-vessel aggregates (Figure 3m-n, Supplementary Fig. 7). Such cellular aggregates were not observed in hiTEV-outlets, in non-treated controls (without LDL, with iMP perfusion = LDL-/iMPs+), nor in native healthy vasculature. However, in atherosclerotic carotid artery explants we identified similar structures (Figure 3m, Supplementary Fig. 8). In LDL-/iMPs+ controls we observed CD11b+ cells adhering to the lumen, invading the tissue, but not colocalizing within plaque-like areas nor accumulating in the inner vessels (Supplementary Fig. 8). Taken together, our findings support the hypothesis that plaque deposition is a process driven by disturbed flow dynamics in combination with and a pro-inflammatory and hyperlipidemic state^5^. Importantly, our multifaceted approach enabled the *in vitro* modeling of plaque buildup within a replica of a human vessel in a reasonable time frame.

**Figure 3.**
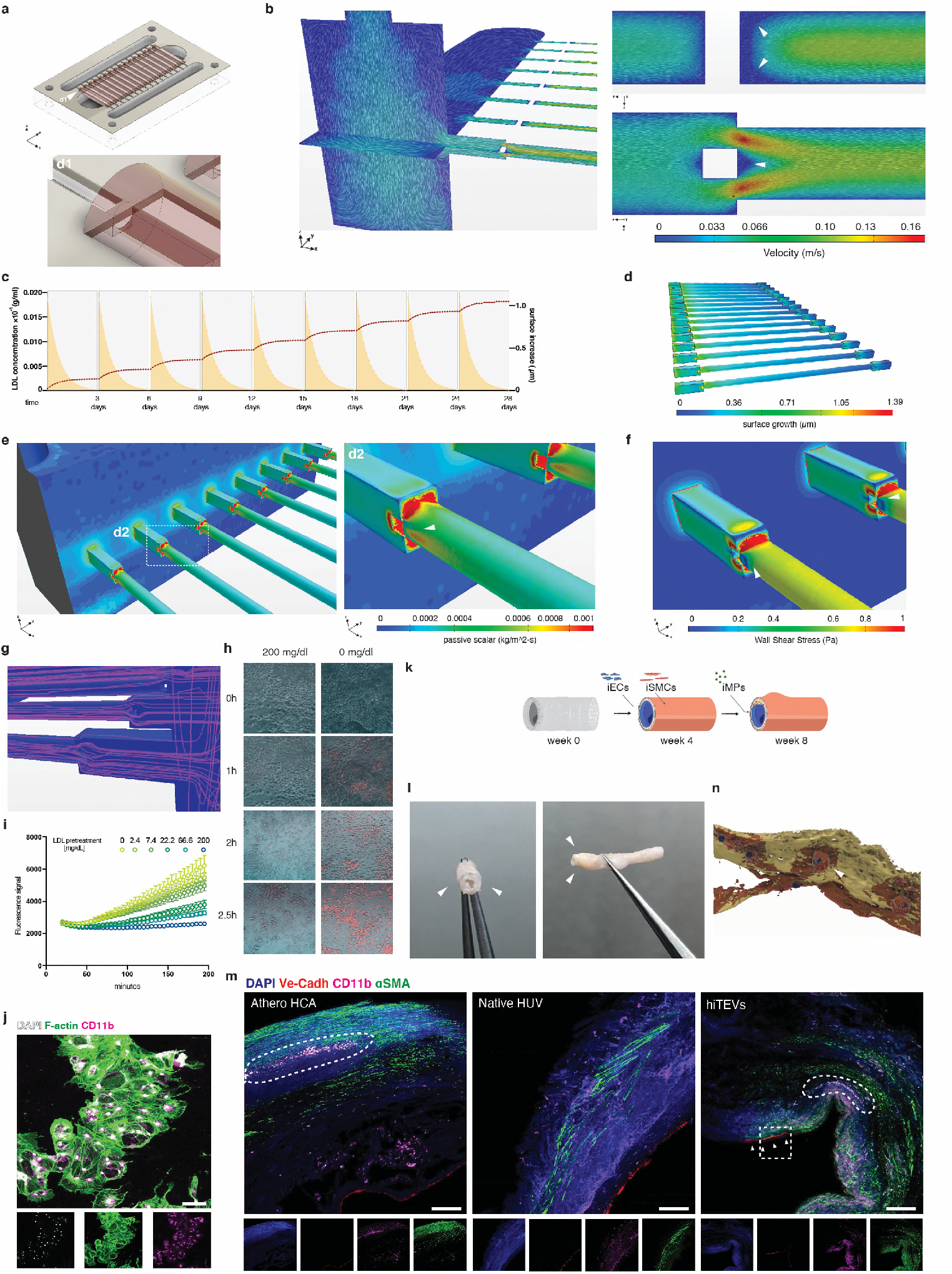
*In silico* and *in vitro* modelling of atherosclerotic plaque deposition. (a) CAD model of the fluidic device loaded with hiTEVs (in red). A detail of a vessel inlet (d1). (b) Flow velocity within the device and at the hiTEV-inlets from different perspectives. Vortexes and whirlpools are indicated by the arrow. (c) CFD prediction of LDL accumulation in the hiTEVs over time (28 days). Concentration of LDL perfused through the device over time (yellow), spiking at every medium change. Prediction of LDL-dependent surface are increase during modeling (red dotted line). The first 1.500 bioreactor cycles upon medium change are reported. (d) Surface growth at the end of the modelling time. (e) Accumulation of LDL (passive scalar) at the inlets and a close-up detail (d2). (f) WSS at the inlets. (g) Modeled hypothetical tracks of the LDL particles within the device (purple lines). (h-i) iMP-macrophages uptake of PhHRodoRed over time. (j) IF characterization of iMP-macrophages. Scale bar 100μm (k) Tissue-engineering and modeling strategy. (l) hiTEVs after 28 days of modeling. Arrows indicate plaque-like regions. (m) IF comparison of atherosclerotic carotid arteries (Athero HCA), Native human umbilical veins (Native HUV), and tissue-engineered arteries from iPSCs (hiTEVs). L = lumen, I = intima layer, M = media layer, S = scaffold remains. Scale bars 200 μm. Round boxes = CD11b+ cell aggregates. Arrows indicate regions where CD11b+ cells adhere to the inner wall of hiTEVs. (n) CAD reconstruction of the highlighted square region in Fig. 3(m) performed from images acquired at the confocal mycroscope. iMPs adhering to hiTEV-lumen (CD11 b+ cell body in dark red, nuclei in blue, Ve-Cadh in light brown).

### 3. Automated clustering identifies atheroma signature cell populations in modeled and native plaques

To validate our model, we compared plaque-resident cell populations from hiTEV-inlets to cells from hiTEV-outlets, as well as cells from native explants deriving from human atherosclerotic and non-atherosclerotic vessels. We applied multiparameter flow-cytometry combined with dimensionality reduction and clustering tools (Figure 4a-c). First, we investigated the changes in cell-cluster abundances between native healthy and native diseased explants. We found that native atherosclerotic lesions had proportionally less monocytes/macrophages populations (cluster 2, cluster 3, p = 0.04), and overall dendritic cells (cluster 5, cluster 6, p = 0.03) when compared with their healthy counterpart. However, native lesion showed proportionally more dendritic cells characterized by low HLA-DR levels (CD11c+, CD31+, HLA-DR^low^), and high surface expression of LDL scavenger receptor (CD11c+, CD36+, HLA-DR^low^) (cluster 8, cluster 7, p = 0.05). Next, we compared the cell population abundances between hiTEV-inlets and hiTEV-outlets. We found that hiTEV-inlets had proportionally fewer monocytes/macrophages (cluster 2, *p* = 0.04), but significantly more dendritic cells with lower levels of HLA-DR, and high surface expression levels of LDL scavenger receptor (Cluster 7, CD11c+/CD36+/HLA-DR^low^) (*p* = 0.05) than hiTEV-outlets (Figure 4a-c, Supplementary Fig. 9). We then looked at the cell populations within healthy native explants and hiTEV-outlets, and we did not find any significant difference in the targeted cell populations between the two. Finally, we compared native plaques and hiTEV-inlets. Here, we found similarities in population abundances of dendritic cells with low HLA-DR levels and high levels of LDL scavenger receptor (cluster 7, ns, *p* = 0.27), SMCs/myofibroblasts (cluster 9, ns, *p* = 0.68) and monocytes/macrophages (cluster 1, ns, *p* = 0.10). However, native plaques still had higher abundance of dendritic cells with low HLA-DR levels (cluster 8, *p* = 0.04) (Figure 4a-c, Supplementary Fig. 9). These results offer an in-depth profile of the signature cell populations within hiTEV-inlets and native plaques, suggesting that our atherosclerosis-on-a-chip model may well approximate reality.

**Figure 4.**
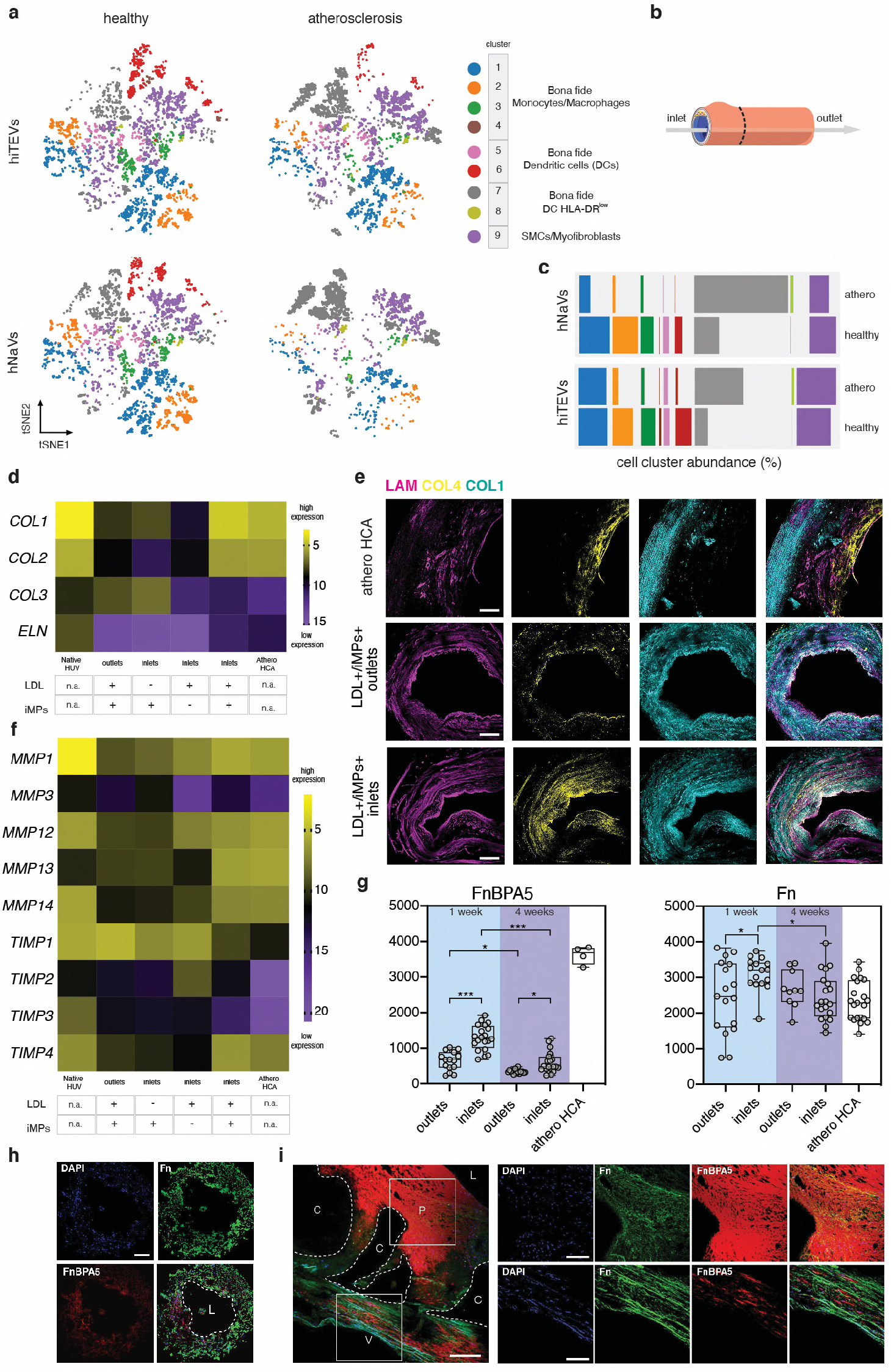
Cellular and extracellular characterization of native and tissue-engineered plaques. (a) tSNE plots with overlaid FlowSOM clustering displaying relative cell population abundances. Cell populations in healthy tissue-engineered hiPSC-derived small-caliber vessels (hiTEV-outlets) and human native healthy vasculature (healthy hNaVs) are compared with their diseased counterpart: hiTEV-inlets and atheroslcerosis hNaVs, respectively. (b) Overview of the analysed hiTEV-regions. (c) Display of relative population abundances in an exploded stacked bar chart. (d) RT-qPCR of Collagens and elastin in hiTEVs and native tissues (Native HUV, Athero HCA). (e) IF analysis of ECM components. LDL+/iMPs+ treated hiTEV-inlets are compared to their internal negative control LDL+/iMPs+ treated hiTEV-outlets. Scale bars 200 μm. (f) RT-qPCR of genes involved in ECM remodeling: matrix metalloproteinases (MMPs) and tissue inhibitors of matrix metalloproteinases (TIMPs). (g) FnBPA5 peptide binding (red) in hiTEV-inlets vs hiTEV-outlets. Counterstaining of fibrillar fibronectin (green) in the same specimens. (h) Distribution of FnBPA5 (red) and fibrillar fibronectin (green) within the ECM of a representative dissected hiTEV-inlet and (i) native carotid plaque. P = atherosclerotic plaque, C = former calcified regions, V= vessel tunica media, L = lumen. Scale bars 200 μm.

### 4. Transcriptional profile of ECM homeostasis, and FnBPA5 peptide-binding patterns provide a signature of increased extracellular matrix remodeling in engineered plaques

To explore the extracellular matrix remodeling landscape of atherosclerotic plaques, we investigated the transcriptional profile of classical ECM-related genes throughout hiTEVs and human samples. We found substantial transcriptional differences in collagen and elastin genes between LDL+/iMP+ hiTEV-inlets and hiTEV-inlets outlets. In detail, hiTEV-inlets had fewer transcripts for *COL1, COL2* and *ELN* (*p* = 0.04, *p* = 0.03, *p* < 0.001 respectively) compared to hiTEV-outlets, hinting to a possible suppressive effect on such ECM-related genes by a combination of iMPs, LDL and the peculiar flow dynamics at the hiTEV-inlets (Figure 4d, Supplementary Table1). Importantly, the microanatomical localization of the ECM components and relative expression of genes for ECM anabolism from hiTEV-inlets and native plaques were similar (*COL1* ns *p* = 0.39, *COL2* ns *p* = 0.42, *COL3* ns *p* = 0.38, *ELN* ns *p* = 0.27) (Figure 4e). Next, we compared the gene expression profile of markers for ECM remodeling, including matrix metalloproteinases (MMPs) and their tissue inhibitors (TIMPs) (Figure 4f, Supplementary Table2, Supplementary Table3). We found that hiTEV-inlets had low *TIMP1* (*p* = 0.03) and high MMP13 (*p* = 0.04) relative transcript abundance compared to hiTEV-outlets, and that LDL alone exerted a downregulation effect on *MMP3* (*p* = 0.03), while upregulating *TIMP2* (*p* = 0.05). When we compared native plaques to hiTEV-inlets, we reported a significant downregulation of *TIMP2* and *TIMP3* (p < 0.001 in both cases). However, no other differences were uncovered, indicating that our model may well recapitulate the ECM transcriptional profile of human atherosclerotic lesions. Last, we focused on fibronectin (Fn) fibers, one of the first assembled and remodeled ECM fibers during inflammation^26,27^ (Figure 4g-i). Based on previous evidence^20,26^, we hypothesized that the peculiar and dynamic flow environment can affect Fn remodeling in both native plaques and in hiTEVs. To test our hypothesis, we stained human native plaques using a fluorescently labeled FnBPA5 peptide, capable of binding to fibronectin fibers with a relaxed conformational state following exposure to low strain. Such low-strain regions are particularly common in fibrotic and overly remodeled tissues^27^. We found that the FnBPA5 peptide selectively binds to plaque regions, indicating a prevalence of Fn fibers under low strain (Figure 4i). Furthermore, we stained hiTEVs and we found that the FnBPA5 peptide could selectively bind to ECM regions at the hiTEV-inlets already after one week (*p* < 0.001) and until the modeling endpoint (*p* = 0.01) (Figure 4g-h). Overall, our data indicate that the athero-prone modeling environment promotes a profound reshuffling of ECM components within the tissues. The observations that human late-lesions and hiTEV-inlets were positive for FnBPA5 staining, support our hypothesis stating that fluid dynamic-dependent mechanical strains are promoters of Fn and, on a broader perspective, of ECM remodeling.

## Discussion

Despite decades of atherosclerosis research, the detailed pathophysiology of plaque seeding, progression and rupture have yet to be elucidated^2^. To dissect each step of the process and to ultimately restrain disease progression, it is pivotal to develop human cell-based disease models with high translational relevance. This work introduces a human plaque-on-a-chip *in vitro* model entirely developed with tissue-engineered vessels from syngeneic iPSC-derived cells. To serve our modeling purpose, we established a novel vascular tissue engineering (VTE) approach which included tailored iPSCs differentiation strategies and the alternation between static and dynamic culturing conditions in a customized fluidic device. Other VTE approaches have attempted to recapitulate the human anatomy using hiPSC-derived or native cells^19,28–30^. However, these works lack sufficient molecular and functional characterization of the employed cellular compartment, detailed histological analyses, and the description of a reproducible culturing setup. Here, we presented a detailed collection of protocols, data and materials to overcome these issues. Next, we developed a CFD analysis that highlighted regions at the hiTEV-inlets characterized by flow disturbances and whirlpools, which were more prone to long-lasting impact and therefore accumulation of LDL particles. Therefore, our CFD model predicted possible atheroprone, hyperplasic, LDL-laden inflammatory regions. These findings are in line with previous reports^10,13,15^ and complement our *in vitro* modeling results from the perfusion of hiTEVs in atheroprone conditions^10,31,32^. Our findings support the hypothesis that atherosclerotic plaque deposition is a process orchestrated by the phagocytic ability of macrophages and dendritic cells, gathering in the sub-intima, and by shear-dependent blood-wall mass transport, providing a first *in vitro* evidence of such combined processes^1,33–37^. We also identified atheroma signature cell populations through immune cell profiling of the human plaque in both native and tissue-engineered specimens, adding a new layer of information to previous discoveries^21,38^. We then dissected and compared the extracellular matrix milieu of native and tissue-engineered plaques finding a close resemblance in transcriptional levels of genes for ECM assembly and remodeling. Importantly, specimens from hiTEV-inlets showed a downregulation of elastin transcripts to levels similar to those found in native plaques and differently from what was observed in healthy native samples. Similar findings have been reported by previous *in vivo* studies^39,40^. Last, we found similarities between native and bioengineered plaques concerning FnBPA5 peptide-binding patterns. FnBPA5 was previously used to assess the tensional state of fibronectin fibers in tumors^27^. Here, we report for the first time the ability of the FnBPA5 peptide to bind both to native and bioengineered plaque regions, providing a possible diagnostic tool and a hint for the development of novel, less invasive, therapies that may locally counteract plaque progression.

Besides the advancements, it is important to mention the limitations of our approach. First, the flow-cytometry characterization and the qPCR analysis performed on hiPSCs-derived iECs, iSMCs, and iMPs revealed that these cells retain a fetal-like phenotype, a common characteristic of most hiPSC-derived cells, which has been extensively described^29^. However, it is important to mention that, contrary to common belief, atherosclerosis is a vascular condition independent of age, affecting both young and elderly subjects. Atherosclerotic plaques have been found in the vascular system of fetuses^41^, children^42^,and young adults^43^. Conversely, severe consequences of atherosclerosis due to plaque rupture or plaque erosion, maintain a higher prevalence in elderly subjects, and this is mainly due to the presence of other comorbidities exacerbating the disease progress^44^. With this in mind, we believe that the impact of the phenotypical cell age on the model output is only limited. As a second limitation, our model investigates intravascular inflammatory events mediated by macrophage precursors and does not include other cells that might interfere with the etiology of the disease (i.e., T and B cells)^45^. The choice to omit those cell types from the modeling setup was driven by the aim to explore a minimalistic disease scenario to test the response-to-retention hypothesis of early atherogenesis^3^, which we later confirmed. Third, our CFD model lacks accurate rendering of the vascular component, herein considered as an inert element. The model could be improved in the future by including more informative boundary conditions based on data from mechanical testing experiments. Moreover, our computational model could be further enhanced by implementing accurate finite element modeling (FEM) and fluid-to-solid interaction (FSI) studies. Such modeling updates shall be based on two-way coupling computations, where LDL particles are influenced by the system (i.e., medium flow) and by possible impacts with other particles within the device. In conclusion, the implementation of CFD approaches in tissue engineering and personalized medicine is now on the rise^46^. Interdisciplinary approaches based on CFD coupled with tissue-engineering will contribute to the development of more accurate TE models, offering a new toolbox to gather insights into the physiologic and pathophysiologic remodeling phenomena observed in atherosclerosis development but not limited to this field.

## Materials and methods

### 1. Cell culture

#### 1.1 Culture of human induced pluripotent stem cells

Two different induced pluripotent stem cell lines derived from skin fibroblasts were used to conduct the tissue engineering experiments included in this work, HPSI0114i-eipl_1 (RRID:CVCL_AE06) and WTB6 (RRID:CVCL_VM30), were purchased from the European Bank for induced pluripotent stem cells (EBiSC). hiPSCs were thawed into 6 well plates (TPP) that were pre-coated for 1h at RT with 10 *μ*g/ml Vitronectin (Stem Cell Technologies) in CellAdhere dilution buffer (Stem Cell Technologies). Each coated plate was prepared fresh before use. iPSCs were thawed in 1 ml of mTeSR medium (Stem Cell Technologies) with the addition of 10 *μ*M Rho kinase inhibitor (ROCKi, Y-27632; Sigma). After 24h, and upon adhesion of the cells to the bottom of the well, the medium was replenished with mTeSR without ROCKi. The medium was refreshed every day after 1 washing step in DPBS (Gibco). The cells were constantly monitored for signs of differentiation such as the formation not-well defined colony edges, or variations in colony shape from the regular circular-oval morphology. The hiPSCs were passaged at about 70% confluence. Here, cells were detached with ReLeSR (Stem Cell Technologies) at 37°C for about 5 min. Next, mTeSR medium was added in a 1:1 ratio, and the detached cell aggregates were carefully transferred to a 50 ml (Falcon) tube using 5 ml pipettes to avoid the formation of single-cell suspensions. Generally, hiPSCs cultures were passaged every 4-7 days using either a 1:4 or a 1:6 split ratio.

#### 1.2 Culture of endothelial cells

Human umbilical vein endothelial cells (HUVECs) were harvested from healthy specimens of umbilical cords donated to the Institute for Regenerative Medicine of the University of Zurich, within the work frame approved by the Kantonale Ethikkommission Zürich (KEK-Stv-21-2006). Umbilical cords were dissected, and the umbilical vein was isolated, washed in PBS (Gibco), and infused with an enzymatic mix of collagenase/dispase following the manufacturer’s instructions (Sigma) for 20 min. At this point, the enzymatic solution was collected in a 50 ml tube (Falcon). Umbilical veins were then washed in PBS and each wash was collected in the same tube with the rinsed enzymatic solution. EGM-2 medium (Lonza) was added in a 1:1 ratio to the cell suspension before a centrifugation step at 300g for 5 min. HUVECs were pelletized, resuspended in EGM-2 medium, and plated onto 0.1% gelatin-coated (Sigma) 6 well plates. The medium was refreshed every 2-3 days, following 2 washes is PBS. HUVECs were sub-cultured at about 80% confluence in a 1:6 sub-culturing ratio. Human brain microvascular endothelial cells (HBMECs) were purchased from ScienCell Research Laboratories cultured, sub-cultured and processed as described for the HUVECs.

#### 1.3 Fibroblasts and myofibroblasts

Human umbilical vein myofibroblasts (HUVMs) were harvested from healthy umbilical cords donated to the Institute for Regenerative Medicine of the University of Zurich, within the work frame approved by the Kantonale Ethikkommission Zürich (KEK-Stv-21-2006). Umbilical cords were dissected, and the umbilical vein was isolated, washed in PBS and cut into pieces of 1mm × 1mm size. Vein wall fragments were then scattered onto 10cm Petri dishses (TPP) and let adhere to the bottom of the plate for 15 min before the addition of DMEM medium, that was prepared as follows: 10% FBS (Gibco), 1% Glutamax (Gibco) and 1% Penn Strep (Sigma) in DMEM medium (Sigma). After about 10 days, myofibroblasts were sprouting from the tissue fragments forming a layer of about 60% confluence. Here, cells were detached from the dish using Trypsin – EDTA solution 0.025% (Sigma), washed in PBS and sub-cultured at a 1:3 ratio. The DMEM medium was refreshed every 2-3 days, after a wash in PBS. HUVMs were passages at 80% confluence. Human aortic fibroblasts (HAFs) were purchased from ScienCell Research Laboratories and processed as described for HUVMs.

### 2. hiPSCs differentiation

#### 2.1 Differentiation of hiPSCs to arterial endothelial cells

The hiPSCs were differentiated to arterial ECs via mesodermal lineage control performed using VEGF and cyclic AMP, following an adapted version of the protocol published by T.Ikuno *et al*.^47^. In detail, hiPSCs colonies were disaggregated to single cells and plated onto freshly prepared thin-coated Matrigel plates (Corning, 1:60) at a density ranging between 60.000-80.000 cells/cm^2^ (Day 0). Cells were cultured for 72h in mTeSR medium enriched with 4ng/ml FGF-B. (Gibco). At this point, the differentiating cells were further coated with a thin layer of Matrigel (1:60) and after 24h, the medium was replaced with RPMI1640 medium (Sigma) enriched with 2mM Glutamax, 1×B27 supplement minus insulin (Gibco), and 125 ng/ml activin-A (PeproTech). After 18h, the medium was replaced with RPMI1640 medium enriched with 2mM Glutamax, 1×B27 supplement minus insulin, 10ng/ml BMP-4 (Sigma), 10 ng/ml FGF-B and Matrigel (1:60). On Day 8, the medium replacement was performed with RPMI1640 enriched with 2mM Glutamax, 1×B27 supplement minus insulin, 1mM 8bromo-cAMP (Stem Cell Technologies) and 100ng/ml VEGF-A (ThermoFisher). On Day 11, the differentiated CD144+ endothelial cells were magnetically sorted on MACS separation columns (Myltenyi Biotec) and seeded at a density of 10.000 cells/cm^2^ on thin-coated, Matrigel dishes in RPMI1640 medium enriched with 2mM Glutamax, 1×B27 supplement minus insulin, 1mM 8bromo-cAMP, 100ng/ml VEGF-A and 10 *μ*M ROCKi. On day 14, the medium was replaced with arterial specification medium prepared as follows: EGM-2 medium with 1mM 8bromo-cAMP. The arterial specification medium was refreshed on day 16 and the resulting arterial endothelial cells were harvested at day 19. iECs were cultured in EGM-2 medium, transferred onto 0.1% gelatin-coated dishes at a seeding density of 10.000 cells/cm^2^, and passaged when their confluence reached about 80%.

#### 2.2 Differentiation of hiPSCs to contractile smooth muscle cells

hiPSCs were differentiated towards contractile smooth muscle cells using an adapted version of the protocol proposed by Yang *et al*.^48^. In detail, hiPSCs were pushed towards the mesodermal lineage by replacing the regular hiPSCs culturing medium mTeSR with mesodermal medium prepared as follows: 5*μ*M CHIR99021(Stem Cell Technologies), 10ng/ml BMP-4, and 2% B27 with insulin (Gibco) in RPMI1640 medium (Day 0). On day 3, the medium was replaced with RPMI1640 enriched with 25 ng/ml VEGF-A, 25 ng/ml FGF-B and 2% B27 minus insulin. The medium was refreshed on day 5. On day 7, the medium was replaced with RPMI1640 enriched with 5ng/mL PDGF-B (Stem Cell Technologies), 2.5 ng/mL TGF-B (PeproTech), and 2% B27 with insulin. A medium refreshment was performed every other day for the following six days. On day 14, the contractile SMCs were selected by culturing the cells in RPMI1640 with 4mM lactate (Sigma). On day 20 the cells were harvested for further characterization or sub-cultured in SMC medium prepared as follows: 5% FBS, 2mM Glutamax, 5ng/mL PDGF-B, 2.5 ng/mL TGF-B in DMEM F12. The iSMCs were passaged at 80% confluence and seeded at a seeding density of 10.000 cells/cm^2^.

#### 2.3 Differentiation of hiPSCs to macrophage precursors

hiPSCs were differentiated to macrophage precursors (iMPs) following the protocol adapted from van Wilgenburg *et al*.^31^. In detail, 4 million hiPSC were seeded into an AggreWell 800 well (Stem Cell Technologies) to generate embryoid bodies (EBs) using E8 medium (Stem Cell Technologies) and fed daily with medium supplemented with 50 ng/mL BMP-4, 50 ng/mL VEGF, and 20 ng/mL SCF (PeproTech). After 4 days, EBs were collected and used to setup macrophage factories in T75 flasks (75 EBs/flask) or T175 flasks (150 EBs/flask) in X-Vivo15 (Lonza), supplemented with 100 ng/mL M-CSF (PeproTech), 25 ng/mL IL-3 (PeproTech), 2 mM Glutamax, 1% penn/strep solution, and 0.050 mM β-mercaptoethanol (Gibco). Fresh medium was added once per week. The iMPs were released into the supernatant after approximately 3 weeks of differentiation and were collected weekly. Upon collection, iMPs were passed through a 40 *μ*m cell strainer (Corning) to obtain a single cell suspension and were either plated at a standard density of 100.000 cells/cm^2^ and differentiated for 7 days to macrophages in X-Vivo15 supplemented with 100 ng/mL M-CSF, 2 mM Glutamax, 1% penn/strep solution for mono-culture assays (i.e., LDL uptake assay), or resuspended in perfusion medium and added to the organ on-a-chip model.

### 3. hiPSCs functional characterization

#### 3.1 Tube formation assay - characterization of iECs

Tube formation assay was performed following an adapted version of a previously described protocol^47^. Specifically, 10.000 differentiated ECs at day 19 were plated on *μ*-Slide Angiogenesis (Ibidi) in 10 *μ*l of Matrigel/well in EGM-2 medium and imaged at time-point 0h, and after 24h. HUVECs and HBMECs were used as controls cell lines. The experiments were conducted on eight cell lines which included iECs_ AE061, iECs_ AE062, iECs_ AE063, from three separate differentiation experiments of the RRID:CVCL_AE06 hiPSCs cell line; iECs_ VM301, iECs_ VM302, iECs_ VM303, from three different differentiation experiments of the RRID:CVCL_VM30 hiPSCs cell line; HUVECs and HBMECs as control endothelial cell lines. Two technical replicates were performed for each cell in each separate experiment.

#### 3.2 Gel contraction test - characterization of iSMCs

SMCs derived from hiPSCs were suspended (2.5×10^5^ cells) in 60*μ*l of an 8 mg/ml fibrinogen solution (Sigma). Hence, the fibrinogen cell suspension was mixed with 60 *μ*l of a 5 U/mL thrombin solution (Sigma) and carefully but quickly deposited in the center of a well of a 24 well plate (TTP) and incubated at 37°C for 10 min in order to allow the complete hydrogel polymerization. The gels were cultured in 1 ml DMEM medium containing 5% FBS, 300 U/ml aprotinin (Sigma) to prevent hydrogel size reduction due to fibrinolysis, and 10 *μ*M ROCKi. Gel sizes were measured at day 0 and at day *3*. HUVMs and HAFs served as controls cell lines. A total of eight cell lines were used for this experiment which included iSMCs_ AE061, iSMCs_ AE062, iSMCs_ AE063, from three separate differentiation experiments of the RRID:CVCL_AE06 hiPSCs cell line; iSMCs_ VM301, iSMCs_ VM302, iSMCs_ VM303, from three different differentiation experiments of the RRID:CVCL_VM30 hiPSCs cell line; HUVMs and HAFs as control cell lines. At least 2 technical replicates were performed for each cell line, at each of the two time-points measured, and in each experiment. Data from different cell lines and time-points were analysed togheter via Two-way ANOVA, mixed-effects model with the Geisser-Greenhouse correction, and Tukey’s multiple comparison test, with individual variances computed for each comparison.

#### 3.3 Wound healing assay - characterization of iSMCs

SMCs derived from hiPSCs (4×10^4^ cells) were resuspended in 140*μ*l of DMEM medium containing 5% FBS and 2 mM Glutamax. The cell suspension was distributed in two silicon cell culture reservoirs of a 24-well plate culture-insert (Ibidi) positioned in the wells of 24 well plates. After 2h, upon full cell adherence to the bottom of the plate, the medium was refreshed. After 24 hours, the medium was further replaced with DMEM medium containing 1% FBS and 2 mM Glutamax, and the silicon inserts were removed to allow cell migration. Plates were imaged with a microplate reader (Tecan, SPARK multimode) and cell migration was reported at 0h and 48h post serum starvation. HUVMs and HAFs served as control cell lines. Sample size and statistics are the same as in *“3.2 Gel contraction test - characterization of iSMCs”*.

#### 3.4 LDL uptake – iMPs characterization

Macrophage precursors derived from hiPSCs SFC840-03-03, SFC854-03-02, SFC856-03-04 (previously characterized^32^), HPSI0114i-eipl_1 and WTB6, were seeded at a density of 100.000 cells/cm^2^ in flat, clear-bottom, black 96-well plates (Corning), and differentiated for 7 days in XVivo-15 medium enriched with 100 ng/ml M-CSF, 2mM Glutamax and 1% penn/strep solution. Labelled low-density lipoprotein, pHRodoRed-LDL (Invitrogen) was applied at a concentration of 2.5 *μ*g/ml in Life Imaging buffer (Gibco). Fluorescent images were acquired with the EVOS M7000 imaging system, supplied with an on-stage incubator unit kept at 37°C. Cells were imaged over 3 hours after pHRodoRed-LDL treatment. Fluorescence intensity was also monitored with fluorescence plate reader (Tecan, Infinite M1000pro), heated to 37°C. Four reads per well were acquired, every 5 min, for a total 3 hours and the mean value was used for further analysis. The fluorescence plate reader was set with an excitation wavelength of 560 nm, 10 nm bandwidth, and with an emission wavelength of 585 nm, 15 nm bandwidth. For LDL uptake inhibition, cells were pre-treated with different concentrations of unlabelled LDL for 30 min (LeeBio) until temporary receptor saturation upon pretreatment with 200 mg/dl LDL. Then the medium was removed and directly replaced with Life Imaging buffer containing pHRodoRed-LDL. At least 2 technical replicates were measured for each cell line, for each unlabeled-LDL pretreatment and untreated controls in each experiment.

## 4. Native tissue specimens

Biopsies of human umbilical veins were obtained from umbilical cords donated for research purposes from healthy subjects in accordance with the ethical protocol approved by the Kantonale Ethikkommission Zürich (KEK-Stv-21-2006). Participants have been informed about the research project before the tissue retrieval and the consent has been be sought from each participant. No compensation was provided to the donor subjects. The sample retrieval was performed during a common clinical procedure therefore, no additional risks for the patient were present. Biopsies of carotid branches were obtained from patients undergoing scheduled carotid endarterectomy and shunting, secondary to vascular stenosis in accordance with the protocol approved by the Ethik Kommission der Universität Witten/Herdecke (Nr.79/2012). Inclusion criteria: age ≥ 18 years old, being scheduled to undergo a carotid endarterectomy. Exclusion criteria: malignancy, liver failure, renal failure (Creatinine >180 ug/ml), hereditary coagulation disorders, anaemia (Hb < 10 g/dl), Rheumatic heart disease, Rheumatoid arthritis.

## 5. RT-qPCR

### 5.1 RNA extraction

For RNA extraction the RNeasy Mini Kit (Qiagen) was used. In case of extraction from adherent cell cultures, the lysis buffer RTL with 10 *μ*l/ml β-mercaptoethanol (Sigma) was directly applied to the culturing plate, the solution was collected in low-bind PCR tubes (Eppendorf) and stored at −80°C until further processing. In case of RNA extraction from native tissues or hiTEVs, the tissues were disrupted in a tissue lyser (TissueLyser II, Qiagen) with 3 mm tungsten beads (Qiagen) in an RTL solution with 10 **μ**l/ml β-mercaptoethanol and the tubes were stored at −80°C. The isolated RNA was measured with a spectrophotometer (NanoDrop 2000, Thermo Scientific) and immediately processed for reverse transcription.

### 5.2 Reverse transcription

The reaction was performed for each sample in a final volume of 20 **μ**l reaction mix containing 1 **μ**g of RNA, 1 × PCR buffer, 5mM MgCl_2_, 10 mM dNTPs, 0.625 **μ**M oligo d(T)_16_, 1,875 **μ**M random hexamers, 20 U RNase inhibitors and 50 U MuLV reverse transcriptase (ThermoFisher). The thermocycler settings were as follows: 25 °C for 10 minutes, 42 °C for 1h, followed by 99 °C for 5 minutes. The resulting cDNA was either processed immediately, stored at 4 °C for a maximum of 48h, or transferred at −20 °C for long term storage.

### 5.3 Quantitative real-time PCR (qPCR)

The reaction was performed for each sample in a final volume of 10 *μ*l reaction mix containing 30 ng of cDNA, a 1:1 mix of froward and reverse primers in a final concertation of 100nM and FAST SYBR green (ThermoFisher). The primers for each tested gene are summarized in the Supplementary Table4. The qPCR was performed on StudioQuant7 (Applied Biosystems) and the amplification program was set as follows: 95° C for 5 min, 40 cycles with 95 °C for 10 seconds, 60 °C for 15 seconds, 72 °C for 20 seconds. The software QuantStudio 6 and 7 Flex Real-Time PCR System (Version 1.0) was used for data acquisition. Gene expression levels were normalized over the two housekeeping genes (HKG): the glyceraldehyde 3-phosphate dehydrogenase (*GAPDH*) and the small subunit 18S ribosomal RNA (*18S*). Three technical replicates were measured for each sample analyzed, and for each gene of interest. The statistical analysis was conducted with ordinary Two-way ANOVA with Dunnett’s multiple comparison test, and individual variances computed for each comparison.

## 6. Immunofluorescence analysis

### 6.1 Fixation and staining

Cells grown in 2D cultures were fixed in 4% cold paraformaldehyde (PFA, Sigma) for 20 minutes at RT. Tissues from native specimens and hiTEVs were embedded unfixed in OCT matrix (Tissue-Tek) and transferred onto dry ice to allow water-soluble glycols and resins to solidify. Hence, the samples were processed in a cryostat (Tissue-Tek) and cut in slices of 5 *μ*m thickness. For the immunofluorescence staining, cells and tissue sections were washed twice in PBS, followed by permeabilization with 0.2% Triton-X 100 (Sigma) in 1× PBS for 10 min RT, and blocked in 1%BSA (Sigma), 0.1% Tween-20 (Sigma) in 1× PBS for 20 min RT. Next, the cells were incubated with the primary antibody for 1h at 37 °C, washed three times in 1× PBS, and stained with secondary antibodies for 1h at 37 °C. The specifications concerning the antibodies used for IF are summarized in the Supplementary Table5. After secondary antibody staining, the samples were washed three times in PBS, counterstained with DAPI 0.2 *μ*g/ml (Sigma) and mounted in Vectaschield (Vector Laboratories).

### 6.2 Confocal microscope imaging and analysis

The stained specimens were analysed with a confocal microscope (Leica TCS SP8) where images were acquired as z-stacks. The acquisition speed was set to 400 Hz, and the picture size to 1024 × 1024 pixels. Each acquired image of the z-stack composite was the result of an average of 10 frames from the same region. The data were then processed using the software Fiji (Version 1.52p Java 1.8.0_172 64-bit). The z-stacks from individual positions within the 2D cell culture or 3D tissue were summarized in one image collecting the maximum intensity voxels from the different focal planes. Each acquired channel was singularly processed as described in the previous sentence and a composite image was created as a merged version of the channels of interest.

## 7. Flow-cytometry

### 7.1 Generation of single cell suspensions from tissue-culture vessels, human tissue and control samples

Native vessels and TEVs were cut into small pieces of about 1mm × 1mm, collected into small 1.5 tubes (Eppendorf) and digested at 1.200 rpm in a tube shaker (Eppendorf) in a solution with 1 mg/ml collagenase/dispase, and 0.5 mg/ml Dnase in 1 × HBSS with Ca^2+^ and Mg^2+^ (Gibco) for 30 min at 37 °C. Single-cell suspensions were filtered through 70 μm nylon meshes (BD Biosciences) and washed in HBSS. Frozen peripheral blood mononuclear cells (PBMCs) isolated from donor blood samples provided by the Zurich Blood Bank (Blutspende Zürich, project Nr.6676) were used as control samples, together with pooled macrophage precursors from HPSI0114i-eipl_1 and WTB6 factories. 2D monocultures of HUVECs and HUVMF were also used as controls and were thawed at 37°C and washed twice in PBS at RT before further processing.

### 7.2 Atheroma signature cell population processing, antibody panel and acquisition

Single-cell suspensions were incubated for 30 min at 4 °C in Zombie Aqua live/dead exclusion dye in HBSS without Mg^2+^ and Ca^2+^ (LIVE/DEAD™ Fixable Aqua Dead Cell Stain Kit, Molecular Probes, Thermo Scientific). Next, cells were washed in FACS buffer prepared with 2% (v/v) heat-inactivated FBS, 5 mM EDTA, 0.01% (v/v) NaN_3_ (Sigma) and resuspended in Fc receptor-blocking antibodies (1:20 from stock in FACS buffer, anti-mouse CD16/32 TruStain fcX, Biolegend). After 5 min incubation, cells were treated with antibody master mix for surface marker staining and incubated for 15 min at 4 °C, followed by a last wash in FACS buffer. For identifying atheroma signature cell populations, we a panel of fluorophore-conjugated antibodies against targeted surface markers (Supplementary Table5). For intracellular staining, cells were fixed and permeabilized according to manufacturer’s instructions (Fixation/Permeabilization and Permeabilization Buffer, eBioscience, Thermo Scientific). Cells were incubated with antibody master mix in 1x Permeabilization buffer for 30 min at 4 °C and incubated with AlexaFluor647 anti-αSMA. Fluorescence-minus-one (FMO) samples were included as negative controls. Cells were acquired on a 16-channel LSR II Fortessa flow cytometer (BD Biosciences).

### 7.3 Data cleaning

Via FlowJo (Version 10.0.8, FLOWJO LLC) software, the forward and side scatter signal of every sample was used to exclude any low-size debris and cell-doublets. Furthermore, Zombie Aqua live/dead exclusion dye-positive dead cells were removed. For follow-up analyses we pre-selected all living cells.

### 7.4 Data normalization, visualization and automated clustering

Living-cell data was normalized via Gaussian Normalization Function in R (gaussNorm, version 3.6.1, R Core Team) implemented in RStudio user interface (version 1.1.447, RStudio, Inc.). Using the viSNE function of the online database Cytobank (https://cytobank.org/cytobank, Cytobank, Beckman Coulter), the multiparameter n-dimensional data was processed by nonlinear dimensionality reduction tSNE algorithm (t-distributed stochastic neighbor embedding algorithm)^49^, in order to visualize cell populations in a two-dimensional space, called viSNE plot (settings: 2500 iterations, 30 perplexity). In order to increase sample size for better visualization in viSNE plots, we concatenated all samples of each experimental group a priori via flowCore (R package version 1.44.2, DOI: 10.18129/B9.bioc.flowCore) and Premessa (R package version 0.2.4, https://github.com/ParkerICI/premessa). Automated, unsupervised cell subpopulation clustering was obtained by FlowSOM (self-organizing maps for flow data) analysis, using the hierarchical consensus clustering method (settings: 10 metaclusters, 10 iterations) of Cytobank (https://cytobank.org/cytobank, Cytobank, Beckman Coulter). The automatically identified clusters were overlaid with their corresponding cell events in the viSNE plots, quantified and exported for statistical analysis.

## 8. Tissue engineering of small-caliber tissue engineered arteries

### 8.1. Tissue engineering, microfluidic device and bioreactor setup

Non-woven polyglycolic acid (PGA) meshes (thickness 1mm; specific gravity 70mg/cm^3^; Cellon) were coated with 1.75% poly-4-hydroxybutyrate (P4HB; TEPHA, Inc., USA) by dipping into a tetrahydrofuran (THF) solution (Sigma) as previously done^50^. After solvent evaporation and overnight drying, the scaffolds were shaped as small conduits of length 1.5 cm with an inner diameter of 500 *μ*m. 48h before cell seeding the constructs were placed into 80% EtOH (Sigma) for 30 min, washed twice in 1 × PBS and incubated in SMC medium (See *Materials and Methods section 2.2*) overnight to facilitate cell adherence. First, we seeded iSMCs onto fully the biodegradable PGA/P4HB tubular-shaped conduits. We used a fibrin hydrogel carrier to maximize cell retention in the scaffold mesh, and we then cultured the seeded constructs using a series of fluid dynamic stimuli including both static and dynamic culture for a total of 2 weeks. In detail, iSMCs were seeded within the PGA tubular scaffolds at a seeding density of 6 × 10^6^ cells/cm^2^ in a seeding volume of about 1.5 ml/vessel. Solutions of 750 *μ*l of fibrinogen (Sigma) (10mg/ml of active protein) and 750 *μ*l thrombin (5U/ml) (Sigma) were prepared, and iSMCs were resuspended in the fibrinogen-thrombin mix and distributed both in the inner lumen and the outer surface. After a waiting time of 10 min, when hydrogel crosslinking occurred, the seeded constructs were moved in iSMC medium enriched with 0.9 mM of l-ascorbic acid-2-phosphate (Sigma) (Day 0). At day 2 the medium was replaced with plain iSMC medium and the constructs were maintained in static culture until day 7. At day 7, tissue engineered constructs were moved into the fluidic devices connected to a bioreactor system and maintained under pulsatile flow (physiological-like flow rate 10 ml/min) to induce a mechanical stimulation of the tissue in culture (dynamic culturing). Second, we coated the inner lumen of the conduits with iECs followed by 2 additional weeks of culture, for a total tissue-engineering period of 1 month. In detail, at day 14, iECs were seeded into the lumen of hiTEVs at the seeding density of 0.5 × 10^6^ cells/cm^2^ in a seeding volume of 100 *μ*l. The vessels were then carefully rotated to allow for even seeding throughout the luminal surface. Henceforth, the hiTEVs were kept in static culture in a culturing medium prepared as follows: ½ iSMCs medium and ½ iECs medium for 7 days, and then moved to dynamic culture at day 21 till day 28, when disease modeling begun. The fluidic device used for dynamic culturing and later for disease modeling was built at the department of physics of the ETH Zurich (ETHZ, Switzerland) through computer numerical control (CNC) guided milling of aluminum and poly methyl methacrylate (PMMA) and silicon (VMQ) and was designed to host the simultaneous culture of 16 hiTEVs. The components of the fluidic device were disassembled, thoroughly cleaned, washed in EthOH 80% overnight and sterilized under UV light for 2h before culturing.

### 8.2. Disease modeling in vitro

At day 28, the hiTEVs were perfused with syngeneic iMPs form 2 weeks-old iMP-factories. The culturing medium was prepared as follows: ⅓ iSMCs medium, ⅓ iECs medium and ⅓ iMPs medium. The medium was changed every 3 days and at each medium change 500.000 iMPs/ml were added alongside with LDL, ApoB100 190 mg/dl (LeeBio) in a final volume of 50 ml. The disease modeling time continued for 28 additional days (4 weeks). At the end of the disease modeling time, hiTEVs could be easily harvested for further processing.

## 9. CFD modeling

### 9.1 CAD modeling and mesh creation

A computer-aided design model (CAD) of the components of the fluidic device was generated using Autodesk Fusion 360. The Standard Tessellation Language file (STL) was exported into STAR-CCM+ for mesh rendering. Briefly, the model was imported into 3D CAD and used to create a CAD imprint operation. The inner volume of the device was extracted via seed point and assigned to the region. To increase mesh resolution all boundaries were split into separate entities to make it easier to control the resolution of each separate group of surfaces. Hence, the obstacles were merged to allow targeted surface control on the mesh resolution. The flow within the device was resolved using flow rate of 10 ml/min (velocity of 0.024 m/s) and a fluid density of 999kg/m3, with a dynamic viscosity of 7.8 × 10-4 Pa•s. To verify the reliability of our model, we resolved a second case, with a set velocity at the inlet of 0.1 m/s.

### 9.2 Atherosclerosis modeling in silico

To model the behaviour of LDL particles within the device we used the Navier-Stokes equations and considered LDL as passive scalar with a diffusion coefficient determined by the Stokes-Einstein equations. The Stokes-Einstein equations aim at modeling the diffusion of particles undergoing Brownian motion in a fluid at uniform temperature, and at relating the particle diameter to their diffusion and hence the Schmidt number (ratio of scalar diffusivity to viscous diffusion). All was discretized using unstructured finite-volume. The mass flow inlet generated a flux of LDL into the device, and subsequently produced downstream fluxes on the vessels surfaces. LDL particles were considered as weakly negatively buoyant and, for modeling purposes, was provided at an inlet concentration of 190 mg/dl (or 1.9 ×10^-3^ g/ml) with a mass of 4.98159 × 10^-21^kg and a density of 480.3029 kg/m^3^. Hence, we performed a one-way coupling where only the flow influences particles and not vice versa. We instructed the code to sample impact and to register and remove the LDL particles when hitting a solid surface (e.g., the inner vessel lumen), to not influence any subsequent impact. We measured that about 1.8% of the overall LDL perfused at the inlet remained within device after each iteration. In our calculations, we considered the flow as a steady state and the occurrence of nine medium changes, one every three days of culture, for a total of twenty-eight days of modeling. We predicted that at every medium change, after about 30 min (1427 cycles) all the perfused LDL would be retained within the device and that every three days of culturing, the inner vessel surface growth would be of about 0.143 *μ*m and at the end of the four weeks of modeling would be of 1.05 *μ*m. To estimate the accumulated particles within the vessels (growth) we measured the flux of the passive scalar. In detail, we multiplied the mass flow rate with time to obtain information on the magnitude of the mass flow (m= kg/m^2^), and the putative surface growth rate. Knowing the density of the particles (ρ) we could then infer the surface growth (Δx = m/ρ). Importantly, the growth rate depends linearly on the concentration at the inlet.

## 10. FnBPA5 peptide treatment

### 10.1 FnBPA5 preparation

The FnBPA5 peptides were commercially synthesized and HPLC-purified (Peptide Specialty Laboratories GmbH, Heidelberg, Germany, or Pichem GmbH, Graz, Austria). A spacer consisting of three glycine and one cysteine residue at the N-terminus of the original peptide sequence from *S. aureus* was introduced. The cysteine was introduced for further modifications with fluorescent dye (Cy5). Lyophilized peptides were dissolved in water (TraceSELECT quality, Sigma) with 10% Dimethylformamide (DMF, Sigma) and stored at −20°C for further usage.

### 10.2 FnBPA5 staining of histological tissue sections and data analysis

For immunohistochemistry freshly dissected arteries were embedded in OTC matrix, frozen on dry-ice, and cut into 20 μm sections. Non-fixed cryo-sectioned tissues were thawed, rinsed with PBS to remove all the soluble components, blocked for 30 min with 5% BSA in PBS and incubated for 60 min with 5μg/ml FnBPA5-Cy5. After one washing step the tissues were fixed in 4% PFA in PBS for 10 min. Samples were blocked in PBS with 5% BSA for 30 min and incubated with polyclonal rabbit anti-fibronectin antibody (Supplementary Table5). The primary antibody solution was removed, and samples were washed before incubation with secondary goat anti-rabbit AlexaFluor546 (Supplementary Table5) antibody solution for 45 min. After a washing step in PBS, samples were mounted using Prolong Antifade Gold with DAPI (Invitrogen). The image was acquired using an Olympus FV1000 confocal microscope equipped with a 10× air or 40× water objective. Whole tissue sections were visualized by stitching together individual fields of view using the grid/collection stitching plugin in Fiji. For image analysis at least 30 images per condition from 3 samples were analyzed for pixel-by-pixel signal intensity of Fn and FnBPA5 and graphed without further normalization. The microscope settings were kept unaltered during image acquisition for comparative analysis of signal intensity.

## 11. Statistics

A detailed description of the sample size, statistical test and correction used is reported at each specific methodological section. Data central tendency is generally indicated as mean, and the variation from the central tendency as standard deviation, unless specified differently. We accounted for inter-donor and inter-cell line variability by setting the false discovery rate to 0.1 (90% confidence interval). All the statistical analyses were performed on GraphPad Prism (Version 9.0, GraphPad Software).

## Acknowledgments

AM was supported by the 3R Research Foundation Switzerland (3R-Project 135-13), by the Start-up grant from the Center for Applied Biology and Molecular Medicine (CABMM) of the University of Zurich, and by a Spark grant from the Swiss National Science Foundation (SNSF, CRSK-3_190579/1). AM acknowledges Stephanie Davaz for the technical help in culturing and characterizing iECs and iSMCs, and Dr. Emanuela Fioretta for the insightful scientific exchanges. WH was supported by a SNSF Sinergia grant (177195) and a pilot grant of the *Oxford* – *McGill* – Zurich Partnership in Neuroscience.

## Author Contributions

AM conceptualized the experiments, designed the model and the microfluidic device components under BW and SPH supervision. AM differentiated and characterized hiPSCs and performed the tissue engineering experiments. CG and AM performed the flow cytometry experiments and analyzed the results. VH and VV provided the FnBPA5 peptide. VH, and AM stained, acquired, and analyzed the samples. KC and JHW developed the CFD model. WH and AM differentiated hiPSCs into iMPs and maintained the iMPs factories for the experiments. WH performed LDL-uptake assays. AVE, BW, AM, and SPH obtained fundings for the experiments. AM wrote the manuscript with the help of CG, WH, VH, and MYE. All the co-authors contributed to the final version of the manuscript.

## Data availability

The authors declare that all data supporting the findings of this study are available within the article and its supplementary material files, or from the corresponding authors upon reasonable request.

