## Supplementary figures and tables for "Human induced pluripotent stem cell-derived vessels as dynamic atherosclerosis model on a chip"

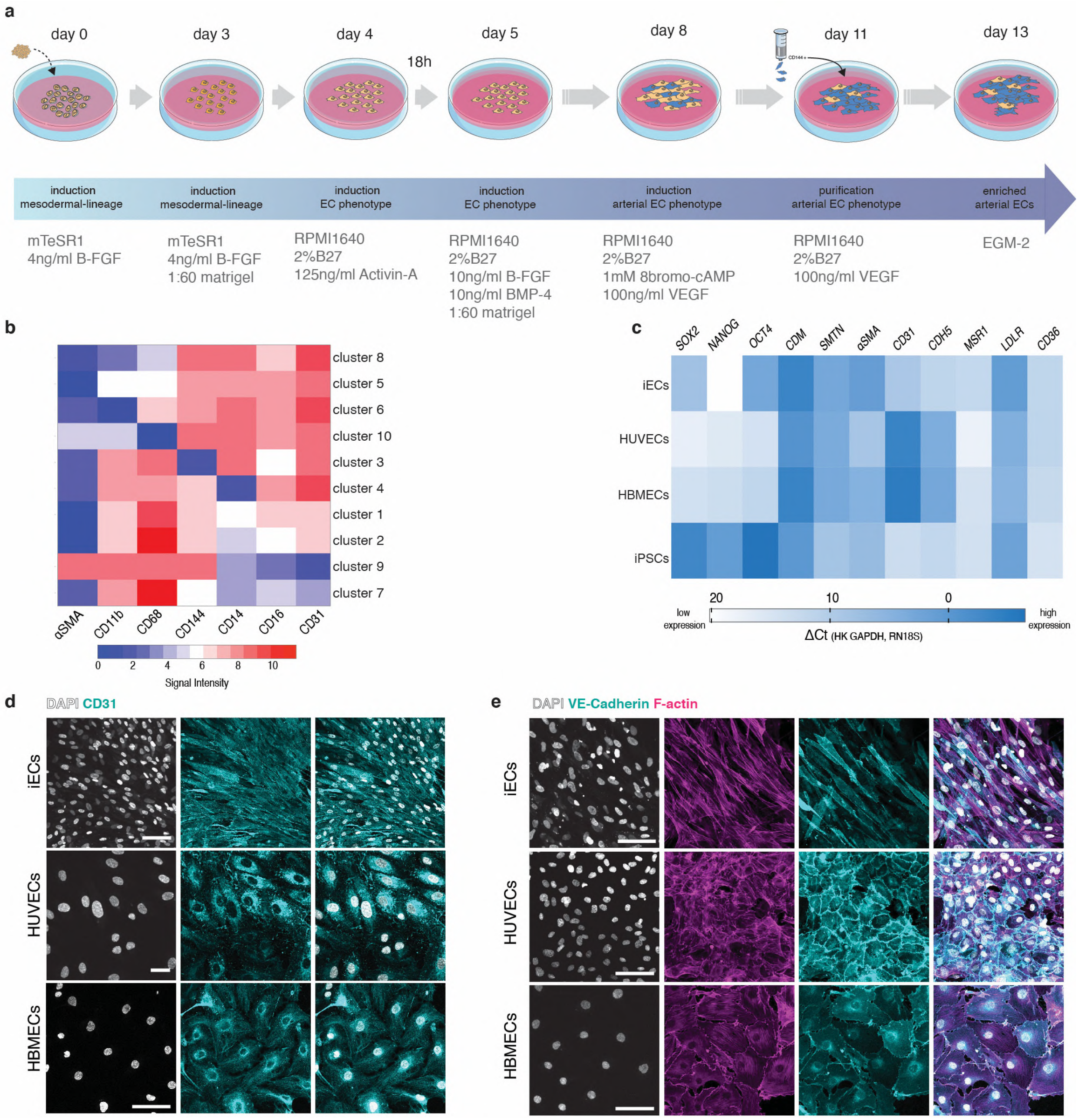


**Supplementary Figure 1. Generation and characterization of arterial endothelial cells from iPSCs.** (a) Differentiation of human iPSCs to arterial endothelial cells at a glance. (b) Fluorescence signal intensity from the flow-cytometry characterization of different endothelial cells (iECs, HUVECs, and HBMECs). (c) RT-qPCR characterization and comparison of different endothelial cells (iECs, HUVECs, and HBMECs). (d) IF characterization of iECs, HUVECs and HBMECs, showing the localization of platelet endothelial cell adhesion molecule precursor (CD31), vascular endothelial cadherin (VE-Cadherin) and F-actin. Scale bars 50µm.

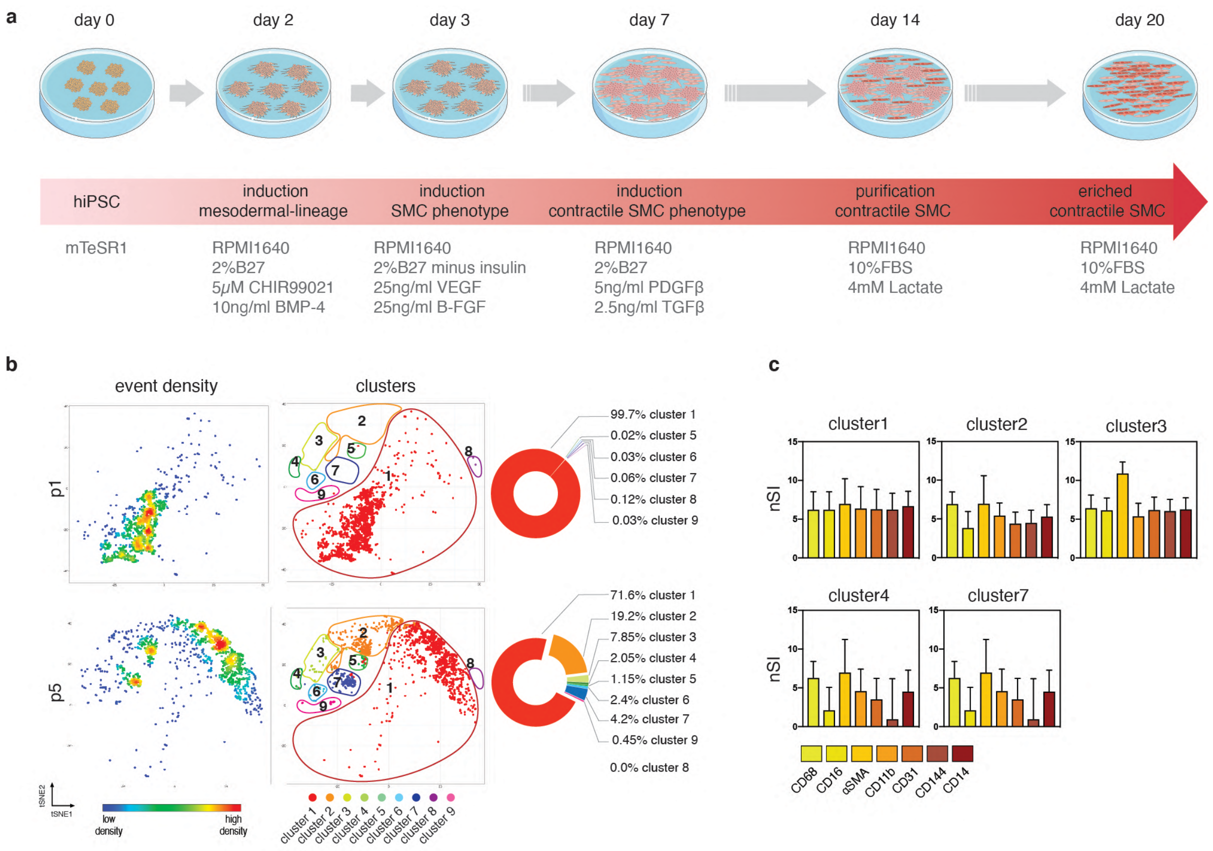


**Supplementary Figure 2. Generation and characterization of contractile smooth muscle cells from iPSCs.** (a) Differentiation of human iPSCs to contractile smooth muscle cells at a glance. (b) flow-cytometry characterization of contractile smooth muscle cells derived from iPSCs. Event density and cluster distribution in the characterized cell populations are shown at two different passages. Passage 1 (p1), the first cell passage once completed the 20days differentiation procedure. Passage 5 (p5), five passages post-differentiation where iSMCs are kept in metabolic medium. P5 iSMCs were used for further experiments (i.e., hiTEV assembly) The relative cluster abundance is shown next to the respective tSNE plots. (c) Normalized median fluorescence signal intensity (nSI) for each cluster and for each tested marker of the multicolor flow-cytometry panel.

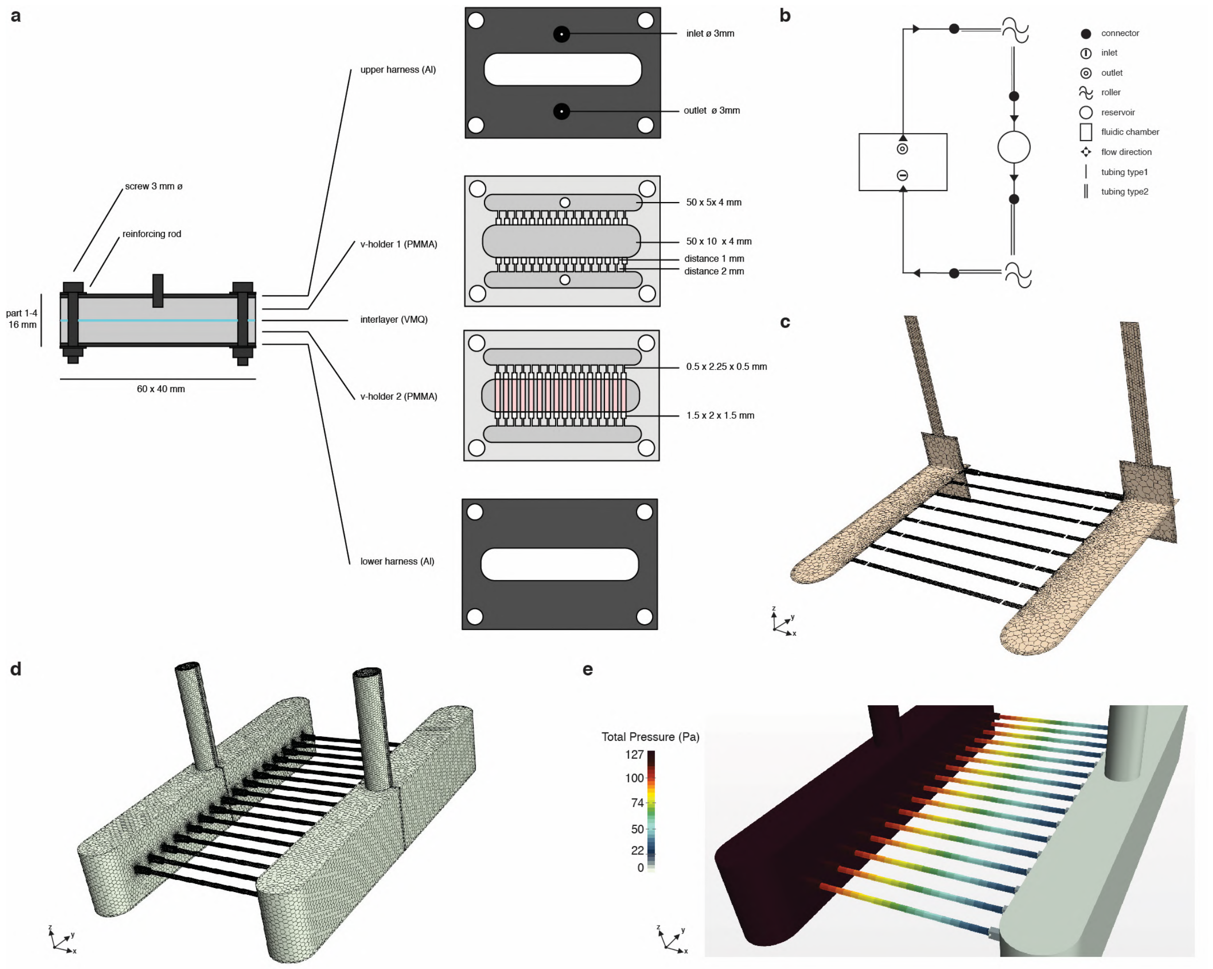


**Supplementary Figure 3. Description of the fluidic device.** (a) Exploded view of the fluidic device. Sizes and components are dscribed. (b) Schematic view of the bioreactor setup. Tubing type 1 = soft clear plastic, Tubing type 2 = opaque reinforced and cyclic strain resistant plastic. (c) Cross-sectional and longitudinal view of the volume mesh of the fluid domain. (d) Overview of the overall volume mesh of the fluid domain. (e) Overview of the total pressure within the fluidic device. The artificial vessels are cultured in a decreasing pressure gradient. hiTEV-inlets at the right side, hiTEV-outlets at the left side.

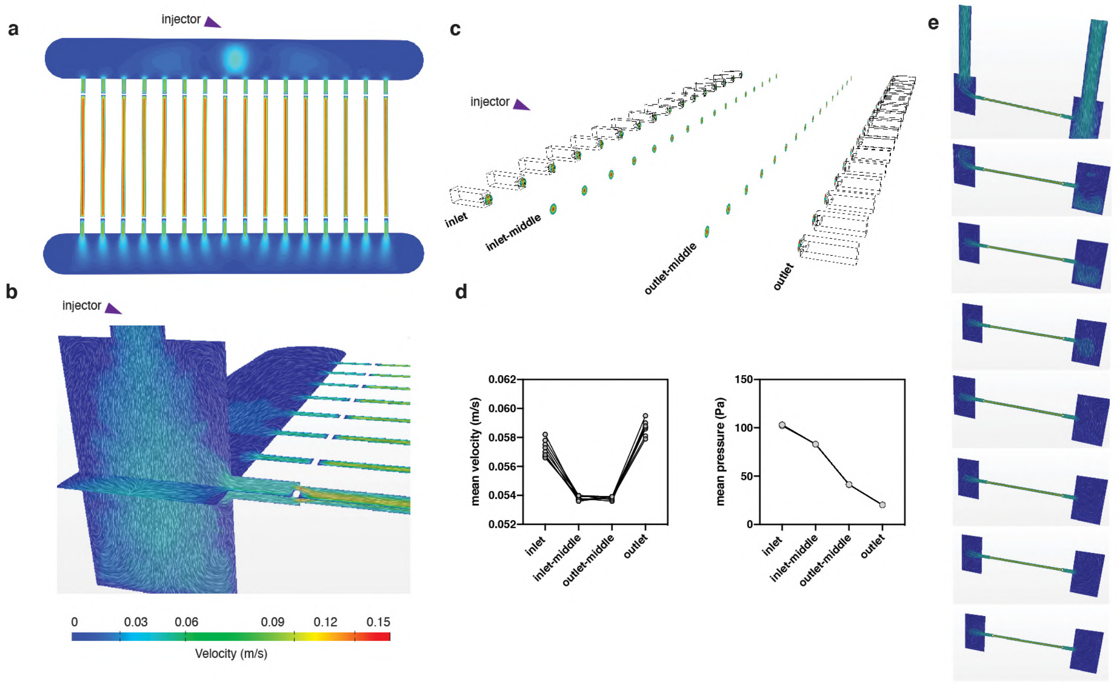


**Supplementary Figure 4. A focus on flow velocity and pressure.** (a) Flow resolution for the velocity vector. (b) A depiction of the flow velocity vectors within the chamber. (c) Flow velocity quantification in four different regions of the tissue-engineered vessels: inlet, inlet-middle region, outlet-middle region, outlet. (d) Quantification of mean flow velocity and pressure changes at different positions within the vessels. (e) Flow velocity in each of the eight vessels representing the flow in half of the chamber. Injector pipe located to the right-side of each panel. Panels are displayed from top to bottom in increasing distance from the injector pipe.

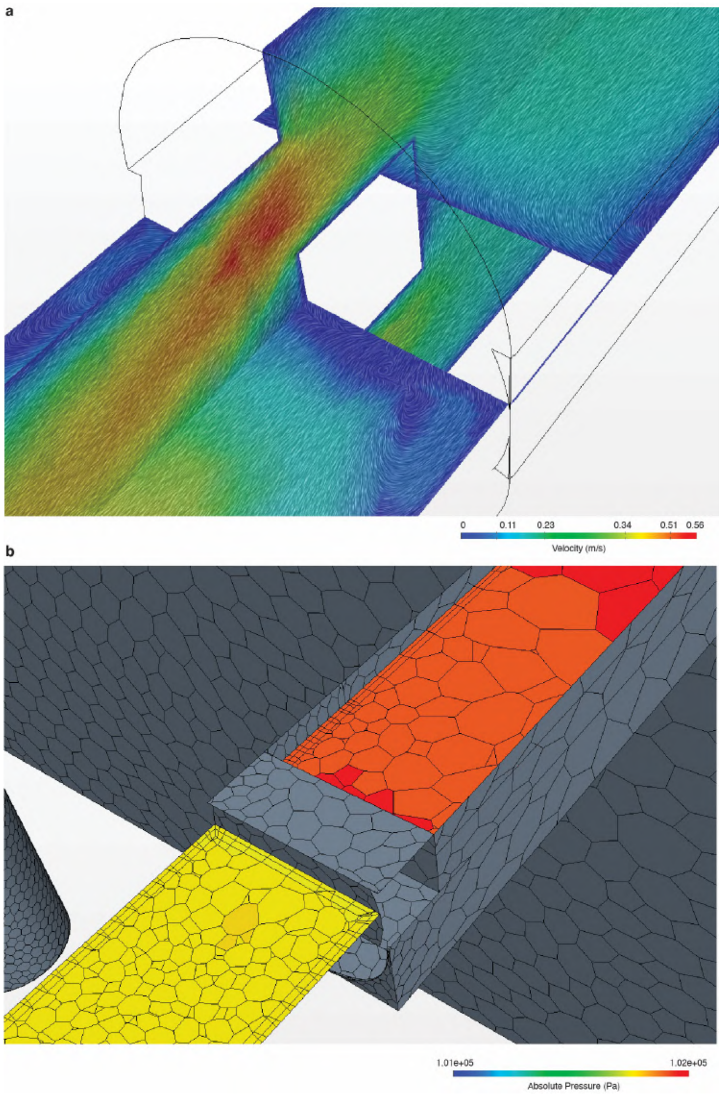


**Supplementary Figure 5. Details of the CFD model.** (a) Velocity field in the 0.1 m/s case. Formation of steady vortexes and whirlpools at the inlet region. (b) Absolute Pressure field superimposed to the mesh domain. Outlet region. 0.1 m/s case.

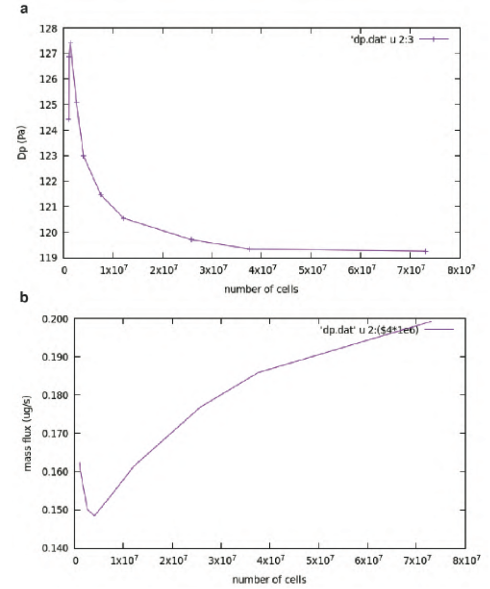


**Supplementary Figure 6. Mesh convergence, a quality control of the CFD model.**

(a) Pressure variation in accordance with the number of computed cells within the mesh.

(b) Mass flux variation in accordance with the number of cells.

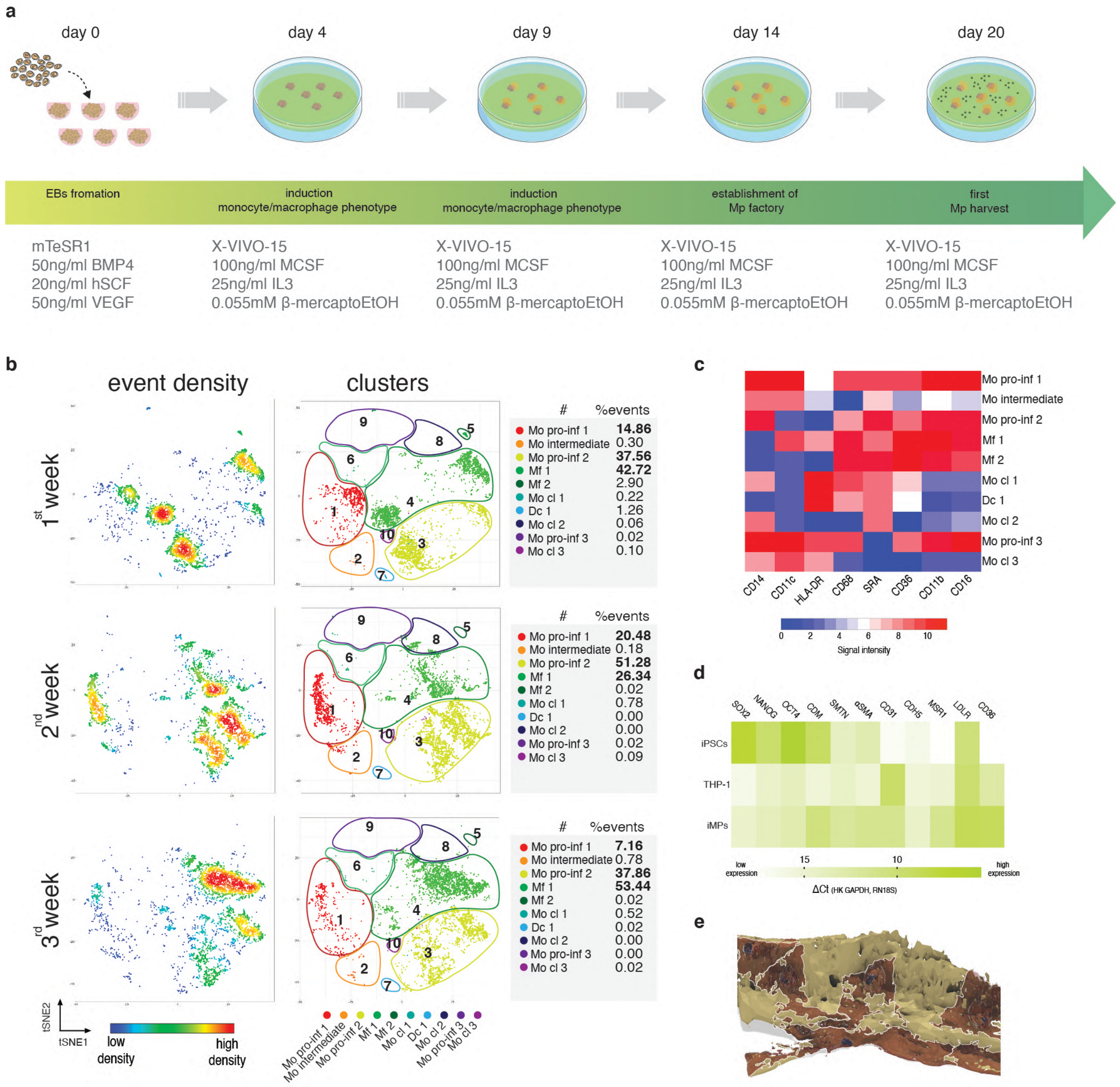


**Supplementary Figure 7. Macrophage precursors form iPSCs - Differentiation, characterization and disease modeling.** (a) Differentiation protocol of hiPSCs into iMPs at a glance. (b) Characterization of iMPs from factories at different culturing time-points. Event density and detailed cluster abundance from multiparameter flow-cytometry analysis. (c) Heat map of surface antigen expression levels from flow-cytometry analysis. (d) RT-qPCR comparing gene expression levels of iPSCs, THP-1 monocytic/macrophage cell line and iMPs from 2-week-old factories. (e) CAD reconstruction from confocal imaging of CD11b+ iMPs adhering to the inner lumen of hiTEVs. The putative perimeter of each cell is highlighted with a white rim.

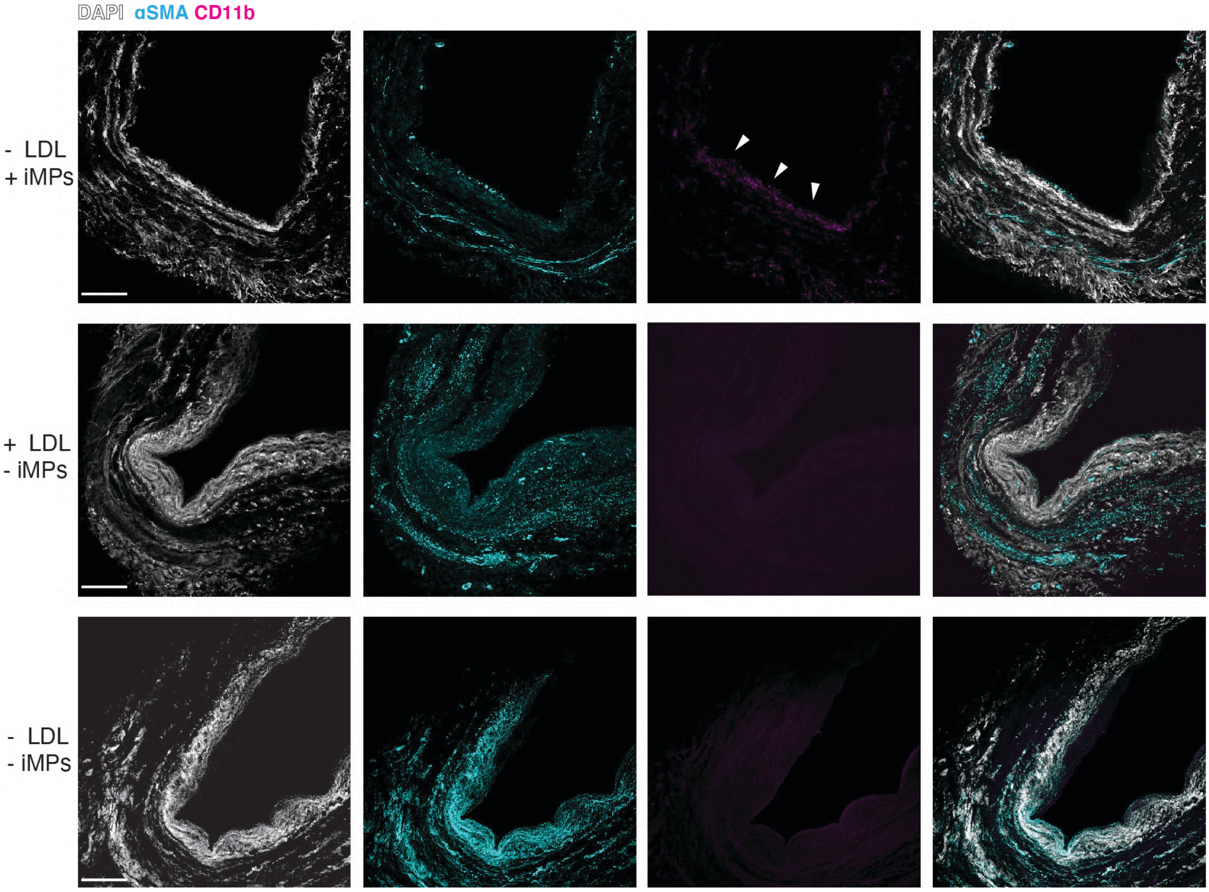


**Supplementary Figure 8. Disease modeling controls.** Immunostaining of control samples from sections collected at the hiTEV-inlets: (1) hiTEVs modeled for 28 days without LDL but with the addition of iMPs. (2) hiTEVs treated with LDL but not perfused with iMPs (3) hiTEVs cultured without adding LDL and iMPs. Scale bars 200 µm. Arrows indicate CD11b+ cells.

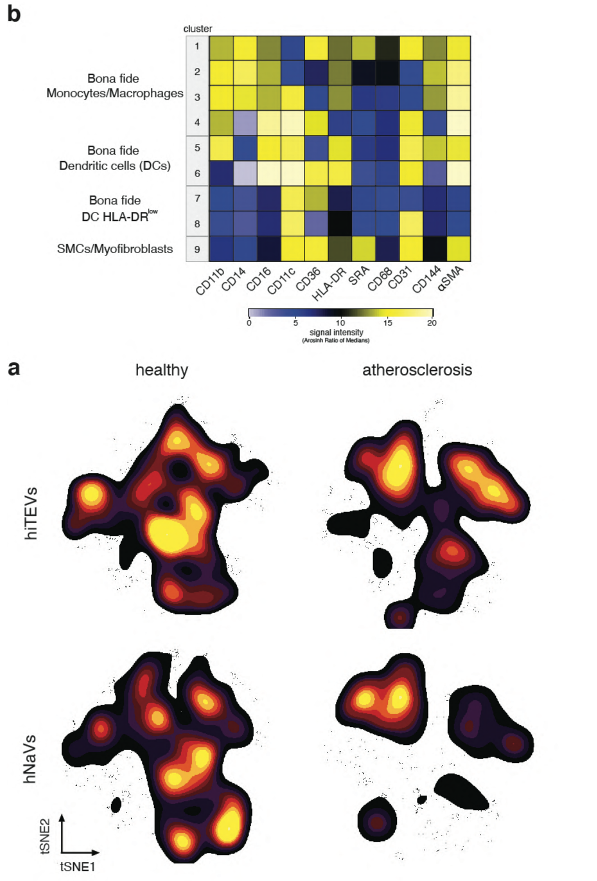


**Supplementary Figure 9. Surface antigen expression heatmap and event density distribution within tSNE.** (a) Heatmap representing the differential median surface expression levels of key lineage markers of intravascular cell populations. (b) event density distribution overlayed to the respective tSNE graph.

| ***COL1*** | VS | Predicted (LS) mean diff. | Discovery? | q value | Individual P Value |
|---|---|---|---|---|---|
| healthy native vessel | carotid plaque | -2.077 | No | 0.12 | 0.14 |
| healthy native vessel | hiTEV LDL+/Mp+ healthy | -5.074 | Yes | 0.007 | 0.003 |
| healthy native vessel | hiTEV LDL+/Mp- | -6.889 | Yes | <0.001 | <0.001 |
| healthy native vessel | hiTEV LDL-/Mp+ | -4.604 | Yes | 0.01 | 0.008 |
| healthy native vessel | hiTEV LDL+/Mp+ plaque | -1.377 | No | 0.28 | 0.42 |
| carotid plaque | hiTEV LDL+/Mp+ healthy | -2.998 | Yes | 0.04 | 0.03 |
| carotid plaque | hiTEV LDL+/Mp- | -4.812 | Yes | 0.003 | <0.001 |
| carotid plaque | hiTEV LDL-/Mp+ | -2.527 | Yes | 0.07 | 0.07 |
| carotid plaque | hiTEV LDL+/Mp+ plaque | 0.6993 | No | 0.39 | 0.62 |
| hiTEV LDL+/Mp+ healthy | hiTEV LDL+/Mp- | -1.815 | No | 0.21 | 0.29 |
| hiTEV LDL+/Mp+ healthy | hiTEV LDL-/Mp+ | 0.4703 | No | 0.46 | 0.78 |
| hiTEV LDL+/Mp+ healthy | hiTEV LDL+/Mp+ plaque | 3.697 | Yes | 0.04 | 0.03 |
| hiTEV LDL+/Mp- | hiTEV LDL-/Mp+ | 2.285 | No | 0.14 | 0.18 |
| hiTEV LDL+/Mp- | hiTEV LDL+/Mp+ plaque | 5.512 | Yes | 0.004 | 0.001 |
| hiTEV LDL-/Mp+ | hiTEV LDL+/Mp+ plaque | 3.227 | Yes | 0.07 | 0.06 |

| ***COL2*** | VS | Predicted (LS) mean diff. | Discovery? | q value | Individual P Value |
|---|---|---|---|---|---|
| healthy native vessel | carotid plaque | -0.4532 | No | 0.38 | 0.74 |
| healthy native vessel | hiTEV LDL+/Mp+ healthy | -3.971 | Yes | 0.02 | 0.02 |
| healthy native vessel | hiTEV LDL+/Mp- | -4.112 | Yes | 0.02 | 0.02 |
| healthy native vessel | hiTEV LDL-/Mp+ | -5.406 | Yes | 0.006 | 0.002 |
| healthy native vessel | hiTEV LDL+/Mp+ plaque | -0.5332 | No | 0.38 | 0.75 |
| carotid plaque | hiTEV LDL+/Mp+ healthy | -3.518 | Yes | 0.02 | 0.01 |
| carotid plaque | hiTEV LDL+/Mp- | -3.659 | Yes | 0.02 | 0.009 |
| carotid plaque | hiTEV LDL-/Mp+ | -4.952 | Yes | 0.003 | <0.001 |
| carotid plaque | hiTEV LDL+/Mp+ plaque | -0.08 | No | 0.42 | 0.95 |
| hiTEV LDL+/Mp+ healthy | hiTEV LDL+/Mp- | -0.1408 | No | 0.42 | 0.93 |
| hiTEV LDL+/Mp+ healthy | hiTEV LDL-/Mp+ | -1.434 | No | 0.26 | 0.4 |
| hiTEV LDL+/Mp+ healthy | hiTEV LDL+/Mp+ plaque | 3.438 | Yes | 0.03 | 0.05 |
| hiTEV LDL+/Mp- | hiTEV LDL-/Mp+ | -1.294 | No | 0.27 | 0.45 |
| hiTEV LDL+/Mp- | hiTEV LDL+/Mp+ plaque | 3.579 | Yes | 0.03 | 0.04 |
| hiTEV LDL-/Mp+ | hiTEV LDL+/Mp+ plaque | 4.872 | Yes | 0.01 | 0.005 |

| ***COL3*** | VS | Predicted (LS) mean diff. | Discovery? | q value | Individual P Value |
|---|---|---|---|---|---|
| healthy native vessel | carotid plaque | -3.544 | Yes | 0.03 | 0.01 |
| healthy native vessel | hiTEV LDL+/Mp+ healthy | 0.7747 | No | 0.49 | 0.65 |
| healthy native vessel | hiTEV LDL+/Mp- | -2.982 | No | 0.1 | 0.08 |
| healthy native vessel | hiTEV LDL-/Mp+ | 1.495 | No | 0.38 | 0.38 |
| healthy native vessel | hiTEV LDL+/Mp+ plaque | -2.429 | No | 0.17 | 0.16 |
| carotid plaque | hiTEV LDL+/Mp+ healthy | 4.319 | Yes | 0.01 | 0.002 |
| carotid plaque | hiTEV LDL+/Mp- | 0.5619 | No | 0.49 | 0.69 |
| carotid plaque | hiTEV LDL-/Mp+ | 5.039 | Yes | 0.004 | <0.001 |
| carotid plaque | hiTEV LDL+/Mp+ plaque | 1.115 | No | 0.38 | 0.42 |
| hiTEV LDL+/Mp+ healthy | hiTEV LDL+/Mp- | -3.757 | Yes | 0.05 | 0.03 |
| hiTEV LDL+/Mp+ healthy | hiTEV LDL-/Mp+ | 0.7203 | No | 0.49 | 0.67 |
| hiTEV LDL+/Mp+ healthy | hiTEV LDL+/Mp+ plaque | -3.204 | Yes | 0.09 | 0.06 |
| hiTEV LDL+/Mp- | hiTEV LDL-/Mp+ | 4.477 | Yes | 0.03 | 0.009 |
| hiTEV LDL+/Mp- | hiTEV LDL+/Mp+ plaque | 0.5528 | No | 0.49 | 0.75 |
| hiTEV LDL-/Mp+ | hiTEV LDL+/Mp+ plaque | -3.924 | Yes | 0.04 | 0.02 |

| ***ELN*** | VS | Predicted (LS) mean diff. | Discovery? | q value | Individual P Value |
|---|---|---|---|---|---|
| healthy native vessel | carotid plaque | -2.915 | Yes | 0.02 | 0.04 |
| healthy native vessel | hiTEV LDL+/Mp+ healthy | -6.214 | Yes | <0.001 | <0.001 |
| healthy native vessel | hiTEV LDL+/Mp- | -7.529 | Yes | <0.001 | <0.001 |
| healthy native vessel | hiTEV LDL-/Mp+ | -6.971 | Yes | <0.001 | <0.001 |
| healthy native vessel | hiTEV LDL+/Mp+ plaque | -3.461 | Yes | 0.02 | 0.04 |
| carotid plaque | hiTEV LDL+/Mp+ healthy | -3.299 | Yes | 0.01 | 0.02 |
| carotid plaque | hiTEV LDL+/Mp- | -4.614 | Yes | 0.002 | 0.001 |
| carotid plaque | hiTEV LDL-/Mp+ | -4.055 | Yes | 0.004 | 0.004 |
| carotid plaque | hiTEV LDL+/Mp+ plaque | -0.5457 | No | 0.27 | 0.69 |
| hiTEV LDL+/Mp+ healthy | hiTEV LDL+/Mp- | -1.315 | No | 0.2 | 0.44 |
| hiTEV LDL+/Mp+ healthy | hiTEV LDL-/Mp+ | -0.7568 | No | 0.27 | 0.66 |
| hiTEV LDL+/Mp+ healthy | hiTEV LDL+/Mp+ plaque | 2.753 | Yes | 0.05 | 0.11 |
| hiTEV LDL+/Mp- | hiTEV LDL-/Mp+ | 0.5585 | No | 0.27 | 0.74 |
| hiTEV LDL+/Mp- | hiTEV LDL+/Mp+ plaque | 4.068 | Yes | 0.01 | 0.02 |
| hiTEV LDL-/Mp+ | hiTEV LDL+/Mp+ plaque | 3.51 | Yes | 0.02 | 0.04 |

**Supplementary Table 1.** Comparison of Collagens and elastin gene expression levels with RT-qPCR.

| ***TIMP1*** | | | Predicted (LS) mean diff. | Discovery? | q value | Individual P Value |
|---|---|---|---|---|---|---|
| healthy native vessel | | carotid plaque | -5.134 | Yes | 0.002 | <0.001 |
| healthy native vessel | | hiTEV LDL+/Mp+ healthy | 0.5273 | No | 0.5 | 0.76 |
| healthy native vessel | | hiTEV LDL+/Mp- | -0.651 | No | 0.5 | 0.7 |
| healthy native vessel | | hiTEV LDL-/Mp+ | -1.247 | No | 0.4 | 0.46 |
| healthy native vessel | | hiTEV LDL+/Mp+ plaque | -3.764 | Yes | 0.05 | 0.03 |
| carotid plaque | | hiTEV LDL+/Mp+ healthy | 5.661 | Yes | <0.001 | <0.001 |
| carotid plaque | | hiTEV LDL+/Mp- | 4.483 | Yes | 0.005 | 0.002 |
| carotid plaque | VS | hiTEV LDL-/Mp+ | 3.887 | Yes | 0.01 | 0.006 |
| carotid plaque | | hiTEV LDL+/Mp+ plaque | 1.37 | No | 0.32 | 0.33 |
| hiTEV LDL+/Mp+ healthy | | hiTEV LDL+/Mp- | -1.178 | No | 0.4 | 0.49 |
| hiTEV LDL+/Mp+ healthy | | hiTEV LDL-/Mp+ | -1.774 | No | 0.32 | 0.3 |
| hiTEV LDL+/Mp+ healthy | | hiTEV LDL+/Mp+ plaque | -4.291 | Yes | 0.03 | 0.01 |
| hiTEV LDL+/Mp- | | hiTEV LDL-/Mp+ | -0.596 | No | 0.5 | 0.73 |
| hiTEV LDL+/Mp- | | hiTEV LDL+/Mp+ plaque | -3.113 | Yes | 0.1 | 0.07 |
| hiTEV LDL-/Mp+ | | hiTEV LDL+/Mp+ plaque | -2.517 | No | 0.17 | 0.14 |

| ***TIMP2*** | | | Predicted (LS) mean diff. | Discovery? | q value | Individual P Value |
|---|---|---|---|---|---|---|
| healthy native vessel | | carotid plaque | -9.737 | Yes | <0.001 | <0.001 |
| healthy native vessel | | hiTEV LDL+/Mp+ healthy | -1.379 | No | 0.33 | 0.42 |
| healthy native vessel | | hiTEV LDL+/Mp- | 2.167 | No | 0.2 | 0.2 |
| healthy native vessel | | hiTEV LDL-/Mp+ | -1.868 | No | 0.24 | 0.27 |
| healthy native vessel | | hiTEV LDL+/Mp+ plaque | -0.8758 | No | 0.41 | 0.61 |
| carotid plaque | | hiTEV LDL+/Mp+ healthy | 8.358 | Yes | <0.001 | <0.001 |
| carotid plaque | | hiTEV LDL+/Mp- | 11.9 | Yes | <0.001 | <0.001 |
| carotid plaque | VS | hiTEV LDL-/Mp+ | 7.869 | Yes | <0.001 | <0.001 |
| carotid plaque | | hiTEV LDL+/Mp+ plaque | 8.861 | Yes | <0.001 | <0.001 |
| hiTEV LDL+/Mp+ healthy | | hiTEV LDL+/Mp- | 3.546 | Yes | 0.05 | 0.04 |
| hiTEV LDL+/Mp+ healthy | | hiTEV LDL-/Mp+ | -0.4893 | No | 0.45 | 0.77 |
| hiTEV LDL+/Mp+ healthy | | hiTEV LDL+/Mp+ plaque | 0.503 | No | 0.45 | 0.77 |
| hiTEV LDL+/Mp- | | hiTEV LDL-/Mp+ | -4.035 | Yes | 0.03 | 0.02 |
| hiTEV LDL+/Mp- | | hiTEV LDL+/Mp+ plaque | -3.043 | Yes | 0.08 | 0.08 |
| hiTEV LDL-/Mp+ | | hiTEV LDL+/Mp+ plaque | 0.9922 | No | 0.41 | 0.56 |

| ***TIMP3*** | | | Predicted (LS) mean diff. | Discovery? | q value | Individual P Value |
|---|---|---|---|---|---|---|
| healthy native vessel | | carotid plaque | -10.97 | Yes | <0.001 | <0.001 |
| healthy native vessel | | hiTEV LDL+/Mp+ healthy | -3.66 | Yes | 0.02 | 0.03 |
| healthy native vessel | | hiTEV LDL+/Mp- | -3.793 | Yes | 0.02 | 0.03 |
| healthy native vessel | | hiTEV LDL-/Mp+ | -4.031 | Yes | 0.02 | 0.02 |
| healthy native vessel | | hiTEV LDL+/Mp+ plaque | -5.985 | Yes | <0.001 | <0.001 |
| carotid plaque | | hiTEV LDL+/Mp+ healthy | 7.314 | Yes | <0.001 | <0.001 |
| carotid plaque | | hiTEV LDL+/Mp- | 7.181 | Yes | <0.001 | <0.001 |
| carotid plaque | VS | hiTEV LDL-/Mp+ | 6.943 | Yes | <0.001 | <0.001 |
| carotid plaque | | hiTEV LDL+/Mp+ plaque | 4.989 | Yes | <0.001 | <0.001 |
| hiTEV LDL+/Mp+ healthy | | hiTEV LDL+/Mp- | -0.1328 | No | 0.41 | 0.94 |
| hiTEV LDL+/Mp+ healthy | | hiTEV LDL-/Mp+ | -0.3713 | No | 0.41 | 0.83 |
| hiTEV LDL+/Mp+ healthy | | hiTEV LDL+/Mp+ plaque | -2.325 | No | 0.11 | 0.17 |
| hiTEV LDL+/Mp- | | hiTEV LDL-/Mp+ | -0.2385 | No | 0.41 | 0.89 |
| hiTEV LDL+/Mp- | | hiTEV LDL+/Mp+ plaque | -2.192 | No | 0.12 | 0.2 |
| hiTEV LDL-/Mp+ | | hiTEV LDL+/Mp+ plaque | -1.954 | No | 0.14 | 0.25 |

| ***TIMP4*** | | | Predicted (LS) mean diff. | Discovery? | q value | Individual P Value |
|---|---|---|---|---|---|---|
| healthy native vessel | | carotid plaque | -0.6019 | No | 0.51 | 0.67 |
| healthy native vessel | | hiTEV LDL+/Mp+ healthy | -3.081 | Yes | 0.08 | 0.07 |
| healthy native vessel | | hiTEV LDL+/Mp- | -4.806 | Yes | 0.02 | 0.005 |
| healthy native vessel | | hiTEV LDL-/Mp+ | -3.649 | Yes | 0.06 | 0.03 |
| healthy native vessel | | hiTEV LDL+/Mp+ plaque | 0.3733 | No | 0.55 | 0.83 |
| carotid plaque | | hiTEV LDL+/Mp+ healthy | -2.479 | Yes | 0.08 | 0.08 |
| carotid plaque | | hiTEV LDL+/Mp- | -4.204 | Yes | 0.01 | 0.003 |
| carotid plaque | VS | hiTEV LDL-/Mp+ | -3.047 | Yes | 0.06 | 0.03 |
| carotid plaque | | hiTEV LDL+/Mp+ plaque | 0.9752 | No | 0.41 | 0.48 |
| hiTEV LDL+/Mp+ healthy | | hiTEV LDL+/Mp- | -1.725 | No | 0.31 | 0.31 |
| hiTEV LDL+/Mp+ healthy | | hiTEV LDL-/Mp+ | -0.5673 | No | 0.52 | 0.74 |
| hiTEV LDL+/Mp+ healthy | | hiTEV LDL+/Mp+ plaque | 3.455 | Yes | 0.06 | 0.04 |
| hiTEV LDL+/Mp- | | hiTEV LDL-/Mp+ | 1.158 | No | 0.41 | 0.5 |
| hiTEV LDL+/Mp- | | hiTEV LDL+/Mp+ plaque | 5.179 | Yes | 0.01 | 0.003 |
| hiTEV LDL-/Mp+ | | hiTEV LDL+/Mp+ plaque | 4.022 | Yes | 0.05 | 0.02 |

**Supplementary Table 2.** Comparison of different tissue inhibitors of matrix metalloproteinases (*TIMP*) gene expression levels with RT-qPCR.

| ***MMP1*** | | | Predicted (LS) mean diff. | Discovery? | q value | Individual P Value |
|---|---|---|---|---|---|---|
| healthy native vessel | | carotid plaque | -4.048 | Yes | 0.01 | 0.004 |
| healthy native vessel | | hiTEV LDL+/Mp+ healthy | -6.94 | Yes | <0.001 | <0.001 |
| healthy native vessel | | hiTEV LDL+/Mp- | -5.021 | Yes | 0.01 | 0.004 |
| healthy native vessel | | hiTEV LDL-/Mp+ | -6.309 | Yes | 0.002 | <0.001 |
| healthy native vessel | | hiTEV LDL+/Mp+ plaque | -3.619 | Yes | 0.08 | 0.04 |
| carotid plaque | | hiTEV LDL+/Mp+ healthy | -2.892 | Yes | 0.08 | 0.04 |
| carotid plaque | | hiTEV LDL+/Mp- | -0.9734 | No | 0.45 | 0.48 |
| carotid plaque | *vs* | hiTEV LDL-/Mp+ | -2.261 | No | 0.16 | 0.11 |
| carotid plaque | | hiTEV LDL+/Mp+ plaque | 0.4288 | No | 0.61 | 0.76 |
| hiTEV LDL+/Mp+ healthy | | hiTEV LDL+/Mp- | 1.919 | No | 0.32 | 0.26 |
| hiTEV LDL+/Mp+ healthy | | hiTEV LDL-/Mp+ | 0.6307 | No | 0.61 | 0.71 |
| hiTEV LDL+/Mp+ healthy | | hiTEV LDL+/Mp+ plaque | 3.321 | Yes | 0.09 | 0.05 |
| hiTEV LDL+/Mp- | | hiTEV LDL-/Mp+ | -1.288 | No | 0.45 | 0.45 |
| hiTEV LDL+/Mp- | | hiTEV LDL+/Mp+ plaque | 1.402 | No | 0.45 | 0.41 |
| hiTEV LDL-/Mp+ | | hiTEV LDL+/Mp+ plaque | 2.69 | No | 0.16 | 0.12 |

| ***MMP3*** | | | Predicted (LS) mean diff. | Discovery? | q value | Individual P Value |
|---|---|---|---|---|---|---|
| healthy native vessel | | carotid plaque | -4.867 | Yes | 0.002 | <0.001 |
| healthy native vessel | | hiTEV LDL+/Mp+ healthy | -2.143 | No | 0.19 | 0.21 |
| healthy native vessel | | hiTEV LDL+/Mp- | -6.254 | Yes | 0.002 | <0.001 |
| healthy native vessel | | hiTEV LDL-/Mp+ | -0.084 | No | 0.64 | 0.96 |
| healthy native vessel | | hiTEV LDL+/Mp+ plaque | -2.193 | No | 0.19 | 0.2 |
| carotid plaque | | hiTEV LDL+/Mp+ healthy | 2.725 | Yes | 0.07 | 0.05 |
| carotid plaque | | hiTEV LDL+/Mp- | -1.386 | No | 0.24 | 0.32 |
| carotid plaque | *vs* | hiTEV LDL-/Mp+ | 4.783 | Yes | 0.002 | <0.001 |
| carotid plaque | | hiTEV LDL+/Mp+ plaque | 2.674 | Yes | 0.07 | 0.06 |
| hiTEV LDL+/Mp+ healthy | | hiTEV LDL+/Mp- | -4.111 | Yes | 0.03 | 0.02 |
| hiTEV LDL+/Mp+ healthy | | hiTEV LDL-/Mp+ | 2.059 | No | 0.19 | 0.23 |
| hiTEV LDL+/Mp+ healthy | | hiTEV LDL+/Mp+ plaque | -0.0505 | No | 0.64 | 0.98 |
| hiTEV LDL+/Mp- | | hiTEV LDL-/Mp+ | 6.17 | Yes | 0.002 | <0.001 |
| hiTEV LDL+/Mp- | | hiTEV LDL+/Mp+ plaque | 4.06 | Yes | 0.03 | 0.02 |
| hiTEV LDL-/Mp+ | | hiTEV LDL+/Mp+ plaque | -2.109 | No | 0.19 | 0.22 |

| ***MMP12*** | | | Predicted (LS) mean diff. | Discovery? | q value | Individual P Value |
|---|---|---|---|---|---|---|
| healthy native vessel | | carotid plaque | 0.01758 | No | >0.99 | 0.99 |
| healthy native vessel | | hiTEV LDL+/Mp+ healthy | -3.337 | No | 0.21 | 0.05 |
| healthy native vessel | | hiTEV LDL+/Mp- | -1.244 | No | 0.77 | 0.47 |
| healthy native vessel | | hiTEV LDL-/Mp+ | -3.513 | No | 0.21 | 0.04 |
| healthy native vessel | | hiTEV LDL+/Mp+ plaque | -0.5257 | No | 0.96 | 0.76 |
| carotid plaque | | hiTEV LDL+/Mp+ healthy | -3.355 | No | 0.14 | 0.02 |
| carotid plaque | | hiTEV LDL+/Mp- | -1.262 | No | 0.67 | 0.36 |
| carotid plaque | *vs* | hiTEV LDL-/Mp+ | -3.53 | No | 0.14 | 0.01 |
| carotid plaque | | hiTEV LDL+/Mp+ plaque | -0.5433 | No | 0.96 | 0.7 |
| hiTEV LDL+/Mp+ healthy | | hiTEV LDL+/Mp- | 2.093 | No | 0.45 | 0.22 |
| hiTEV LDL+/Mp+ healthy | | hiTEV LDL-/Mp+ | -0.1753 | No | >0.99 | 0.92 |
| hiTEV LDL+/Mp+ healthy | | hiTEV LDL+/Mp+ plaque | 2.812 | No | 0.28 | 0.1 |
| hiTEV LDL+/Mp- | | hiTEV LDL-/Mp+ | -2.269 | No | 0.43 | 0.18 |
| hiTEV LDL+/Mp- | | hiTEV LDL+/Mp+ plaque | 0.7183 | No | 0.96 | 0.67 |
| hiTEV LDL-/Mp+ | | hiTEV LDL+/Mp+ plaque | 2.987 | No | 0.27 | 0.08 |

| ***MMP13*** | | | Predicted (LS) mean diff. | Discovery? | q value | Individual P Value |
|---|---|---|---|---|---|---|
| healthy native vessel | | carotid plaque | 4.489 | Yes | 0.006 | 0.002 |
| healthy native vessel | | hiTEV LDL+/Mp+ healthy | 0.7598 | No | 0.42 | 0.66 |
| healthy native vessel | | hiTEV LDL+/Mp- | -0.3575 | No | 0.49 | 0.83 |
| healthy native vessel | | hiTEV LDL-/Mp+ | 0.875 | No | 0.42 | 0.61 |
| healthy native vessel | | hiTEV LDL+/Mp+ plaque | 4.314 | Yes | 0.02 | 0.01 |
| carotid plaque | | hiTEV LDL+/Mp+ healthy | -3.729 | Yes | 0.02 | 0.008 |
| carotid plaque | | hiTEV LDL+/Mp- | -4.846 | Yes | 0.005 | <0.001 |
| carotid plaque | *vs* | hiTEV LDL-/Mp+ | -3.614 | Yes | 0.02 | 0.01 |
| carotid plaque | | hiTEV LDL+/Mp+ plaque | -0.1752 | No | 0.49 | 0.9 |
| hiTEV LDL+/Mp+ healthy | | hiTEV LDL+/Mp- | -1.117 | No | 0.39 | 0.51 |
| hiTEV LDL+/Mp+ healthy | | hiTEV LDL-/Mp+ | 0.1152 | No | 0.49 | 0.95 |
| hiTEV LDL+/Mp+ healthy | | hiTEV LDL+/Mp+ plaque | 3.554 | Yes | 0.04 | 0.04 |
| hiTEV LDL+/Mp- | | hiTEV LDL-/Mp+ | 1.233 | No | 0.39 | 0.47 |
| hiTEV LDL+/Mp- | | hiTEV LDL+/Mp+ plaque | 4.671 | Yes | 0.02 | 0.007 |
| hiTEV LDL-/Mp+ | | hiTEV LDL+/Mp+ plaque | 3.439 | Yes | 0.04 | 0.05 |

| ***MMP14*** | | | Predicted (LS) mean diff. | Discovery? | q value | Individual P Value |
|---|---|---|---|---|---|---|
| healthy native vessel | | carotid plaque | -1.094 | No | 0.4 | 0.43 |
| healthy native vessel | | hiTEV LDL+/Mp+ healthy | -4.877 | Yes | 0.02 | 0.005 |
| healthy native vessel | | hiTEV LDL+/Mp- | -3.619 | Yes | 0.06 | 0.04 |
| healthy native vessel | | hiTEV LDL-/Mp+ | -4.633 | Yes | 0.02 | 0.007 |
| healthy native vessel | | hiTEV LDL+/Mp+ plaque | -1.197 | No | 0.4 | 0.48 |
| carotid plaque | | hiTEV LDL+/Mp+ healthy | -3.783 | Yes | 0.02 | 0.007 |
| carotid plaque | | hiTEV LDL+/Mp- | -2.525 | Yes | 0.09 | 0.07 |
| carotid plaque | *vs* | hiTEV LDL-/Mp+ | -3.539 | Yes | 0.03 | 0.01 |
| carotid plaque | | hiTEV LDL+/Mp+ plaque | -0.103 | No | 0.62 | 0.94 |
| hiTEV LDL+/Mp+ healthy | | hiTEV LDL+/Mp- | 1.258 | No | 0.4 | 0.46 |
| hiTEV LDL+/Mp+ healthy | | hiTEV LDL-/Mp+ | 0.2433 | No | 0.62 | 0.89 |
| hiTEV LDL+/Mp+ healthy | | hiTEV LDL+/Mp+ plaque | 3.679 | Yes | 0.06 | 0.03 |
| hiTEV LDL+/Mp- | | hiTEV LDL-/Mp+ | -1.014 | No | 0.42 | 0.55 |
| hiTEV LDL+/Mp- | | hiTEV LDL+/Mp+ plaque | 2.422 | No | 0.17 | 0.16 |
| hiTEV LDL-/Mp+ | | hiTEV LDL+/Mp+ plaque | 3.436 | Yes | 0.06 | 0.05 |

**Supplementary Table 3.** Comparison of different matrix metalloproteinases (*MMP*) gene expression levels with RT-qPCR.

| | NCBI Gene ID | Location | FW primer (5'→3') | RV primer (5'→ 3') |
|---|---|---|---|---|
| **macrophages and precursors** | | | | |
| *CD36* | 948 | Chromosome 7, NC_000007.14 | TCTTTCCTGCAGCCCAATG | AGCCTCTGTTCCAACTGATAGTGA |
| *LDLR* | 3949 | Chromosome 19, NC_000019.10 | AACTGCCATTGTCTGCTTTA | ACATACCCATCAACGACAAG |
| *MSR1* | 4481 | Chromosome 8, NC_000008.11 | GCAGTGGGATCACTTTCACAA | AGCTGTCATTGAGCGAGCATC |
| **endothelial cells** | | | | |
| *CDH5* | 1003 | Chromosome 16, NC_000016.10 | CCTGGGACTCCACCTACAGA | CCAGAAACGGAGGCCTGATG |
| *CD31* | 5175 | Chromosome 17, NC_000017.11 | AACAGTGTTGACATGAAGAGCC | TGTAAAACAGCACGTCATCCT |
| **smooth muscle cells, myofibroblasts** | | | | |
| *αSMA* | 58 | Chromosome 1, NC_000001.11 | AAAAGACAGCTACTGGGGTGA | GCCATGTTCTATCGGGTACTTC |
| *SMTN* | 6525 | Chromosome 22, NC_000022.11 | TCACCAGAAAGGAACCGACG | CTGTGACCTCCAGCAGCTT |
| *CDM* | 800 | Chromosome 7, NC_000007.14 | GTGCAGAAAAGCAGTGGTGTC | CCGGCTTTGTAGGTTTTGCG |
| **hiPSCs** | | | | |
| *OCT4* | 5460 | Chromosome 6, NC_000006.12 | TCGAGAACCGAGTGAGAGG | GAACCACACTCGGACCACA |
| *NANOG* | 79923 | Chromosome 12, NC_000012.12 | GAACTCTCCAACATCCTGAACCTC | CCTTCTGCGTCACACCATTGC |
| *SOX2* | 6657 | Chromosome 3, NC_000003.12 | GTTGTCAAGGCAGAGAAGAG | GAGAGAGGCAAACTGGAATC |
| **ECM assembly** | | | | |
| *COL1* | 1277 | Chromosome 17, NC_000017.11 | CCGAGACGTGTGGCAAACCCGA | GGCAGTTCTTGGTCTCGTCACAGA |
| *COL2* | 1280 | Chromosome 12, NC_000012.12 | | |
| *COL3* | 1281 | Chromosome 2, NC_000002.12 | GGAGCTGGCTACTTCTCGC | GGGAACATCCTCCTTCAACAG |
| *ELN* | 2006 | Chromosome 7, NC_000007.14 | GCAGGAGTTAAGCCCAAGG | TGTAGGGCAGTCCATAGCCA |
| **ECM remodeling** | | | | |
| *MMP1* | 4312 | Chromosome 11, NC_000011.10 | AAAGGG AATAAGTACTGGGC | CAGTGTTTTCCTCAGAAAGAG |
| *MMP3* | 4314 | Chromosome 11, NC_000011.10 | | |
| *MMP12* | 4321 | Chromosome 11, NC_000011.10 | CATGAACCGTGAGGATGTTGA | GCATGGGCTAGGATTCCACC |
| *MMP13* | 4322 | Chromosome 11, NC_000011.10 | CGTATTGTTCGCGTCATGCCAG | TCTTCCCCTACCCCGCACTTCT |
| *MMP14* | 4323 | Chromosome 14, NC_000014.9 | CCCAGCCCACCCATTGAAGTCT | CCCGACATCCCTCTCCTCTTGC |
| *TIMP1* | 7076 | Chromosome X, NC_000023.11 | TGGAAAACTGCAGGATGGACTCTTG | CAGGGGATGGATAAACAGGGAAACA |
| *TIMP2* | 7077 | Chromosome 17, NC_000017.11 | AAGCGGTCAGTGAGAAGGAAG | GGGGCCGTGTAGATAAACTCTAT |
| *TIMP3* | 7078 | Chromosome 22, NC_000022.11 | CATGTGCAGTACATCCATACGG | CATCATAGACGCGACCTGTCA |
| *TIMP4* | 7079 | Chromosome 3, NC_000003.12 | GCTGCCAAATCACCACCTGCTAC | GTGCCGTCAACATGCTTCATACAGA |
| **Housekeeping** | | | | |
| *18S* | 106631781 | Chromosome 21, NC_000021.9 | CCCGGGGAGGTAGTCACGAAAAATA | CCCGCTCCCAAGATCCAACTAC |
| *GAPDH* | 2597 | Chromosome 12, NC_000012.12 | ACAACTTTGGTATCGTGGAAGG | GCCATCACGCCACAGTTTC |

**Supplementary Table 4.** Primers used for RT-qPCR analyses.

| **IF primary antibodies** | working dilution | host and [clone] | company |
|---|---|---|---|
| CD31 | 1:100 | mouse monoclonal [JC/70A] | abcam (ab9498) |
| vWF | 1:200 | rabbit polyclonal | abcam (ab6994) |
| SM22a | 1:100 | goat polyclonal | abcam (ab10135) |
| SMMHC | 1:100 | rabbit monoclonal [MYH11/2303R] | LSBio (LS-B16305) |
| aSMA | 1:100 | mouse monoclonal [1A4] | abcam (ab7817) |
| Ve-Cadh | 1:250 | goat polyclonal | Santa Cruz Biotechnology (SC-6458) |
| Fibronectin | 1:50 | rabbit polyclonal | abcam (ab23750) |

| **Secondary antibodies/reagents** | working dilution | host and [clone] | company |
|---|---|---|---|
| anti-goat Alexa Fluor 488 | 1:500 | donkey polyclonal | Invitrogen (A32814) |
| anti-mouse Alexa Fluor 488 | 1:500 | donkey polyclonal | Jackson Immuno Research (715-545-150) |
| anti-rabbit Alexa Fluor 488 | 1:500 | goat polyclonal | Invitrogen (A11008) |
| anti-human Collagen 1 Alexa Fluor 488 | 1:200 | rabbit polyclonal | Bioss (bs-10423R-A488) |
| anti-rabbit Alexa Fluor 546 | 1:200 | goat polyclonal | Invitrogen (A11035) |
| anti-mouse Alexa Fluor 647 | 1:500 | rabbit polyclonal | invitrogen (A21239) |
| anti-human CD11b Alexa Fluor 594 | 1:100 | mouse monoclonal [ICRF44] | Biolegend (301340) |
| anti-human Collagen IV Alexa Fluor 594 | 1:200 | mouse polyclonal | Bioss (bs-4595R-A594) |
| anti-human F-actin Phalloidin-Alexa Fluor 633 | 1:500 | - | Invitrogen (A22284) |
| anti-human eNOS AlexaFluor 647 | 1:100 | mouse monoclonal [33/eNOS] | BD Phosflow™ (560102) |
| anti-human laminin Alexa Fluor 647 | 1:300 | rabbit polyclonal | Novus Biologicals (NB300-144AF647) |

| **Flow cytometry antibody panel** | working dilution | host and [clone] | company |
|---|---|---|---|
| anti-human CD31 PacificBlue | 1:100 | mouse monoclonal [WM59] | Biolegend (303114) |
| anti-human VECADH PE-Cy7 | 1:100 | mouse monoclonal [BV9] | Biolegend (348516) |
| anti-human CD14 PerCP | 1:100 | mouse monoclonal [M5E2] | Biolegend (301847) |
| anti-human CD16 Alexa Fluor 700 | 1:125 | mouse monoclonal [3G8] | Biolegend (302025) |
| anti-human CD68 Alexa Fluor 488 | 1:100 | mouse monoclonal [Y1/82A] | Biolegend (333811) |
| anti-human CD11c PE-Cy5 | 1:150 | mouse monoclonal [3.9] | Biolegend (301609) |
| anti-human HLA-DR APC-Cy7 | 1:100 | mouse monoclonal [Y1/82A] | Biolegend (307617) |
| anti-human CD11b Alexa Fluor 594 | 1:250 | mouse monoclonal [ICRF44] | Biolegend (301340) |
| anti-human CD36 Brilliant Violet 605 | 1:150 | mouse monoclonal [CB38, BD] | Fisher Scientific (BDB563518) |
| anti-human SRA PE | 1:150 | mouse monoclonal [7C9C20] | Biolegend (371903) |

**Supplementary Table 5.** Primary antibodies, secondary antibodies, and reagents used for the immunostaining experiments. Detailed flow cytometry antibody panel.
